# The genetic architecture of cerebellar lobules: Insights from the UK Biobank

**DOI:** 10.1101/2022.10.21.513204

**Authors:** Amaia Carriόn-Castillo, Cedric Boeckx

## Abstract

In this work we take advantage of opportunities afforded by the UK Biobank, and complement recent studies examining the genetics of cerebellar volume from that vantage point. We examine the genetic underpinnings of the different cerebellar lob(ul)es, possible reflexes of their evolutionary history, and their genetic relation to psychiatric disorders, cognitive performance and the cortical language network as well as to subcortical regions. Overall, our results show that the cerebellum is a heritable structure, not only globally but also at the regional level. However, our analysis also reveals significant variability across different substructures, justifying the need for a more detailed analysis affording greater structural resolution. Aspects of the “neo-cerebellum”, especially lobule VI/Crus I and, to a lesser extent, Crus II, stand out in our analysis.

## 1 Introduction

There is ever-increasing evidence for the role of the cerebellum in cognitive processes, extending its functions beyond its classical implication in movement coordination and balance [1, 2]. Although the cerebral neocortex continues to be treated as the seat of our “higher-order” cognitive abilities, it is clear that the cerebellum also plays a key role, likely due to its extensive connections with virtually every part of the neocortex (and beyond) [3], as well as its actual surface area [4].

Like the cerebral neo-cortex, the cerebellum consists of multiple areas that exhibit a complex range of connections with other brain regions [5, 6], and is implicated in a wide range of processes, such as sensorimotor, cognitive and social/affective [7, 8, 9, 10, 11]. While the sensorimotor cerebellum is represented mostly in the anterior cerebellar lobe (lobules I to V), with lesions of these areas leading to motor disorders (e.g. cerebellar motor syndrome of ataxia, dysmetria), the posterior portions of the cerebellum (lobules VI-IX, Crus I and Crus II) have been implicated in ‘higher’ cognitive processes, with lesions resulting in the cerebellar cognitive affective syndrome [12]. Consequently, the cerebellum (especially the phylogenetically more recent, posterior lobes) has been implicated in a wide range of psychiatric disorders [13, 14]. There is also mounting evidence for the evolutionary relevance of cerebellar expansion in the hominin lineage [15, 16, 17, 18, 19].

Two recent studies have investigated the genetics of cerebellar volume and have established that it is a heritable structure, identified multiple associated genetic loci, and revealed genetic links with mental disorders [20, 21]. However, these studies focused primarily on the genetic architecture of overall cerebellar volume. We decided to make use of opportunities afforded by the UK Biobank, a unique resource that enables researchers to examine genetic effects on human brain development and disease [22, 23, 24], to further dissect possible genetic contributions to the different cerebellar regions. Thus, in this work we take a complementary approach and utilize genetic analyses to examine cerebellar implication in cognitive processes. To this aim, we investigate the genetic underpinnings of the different cerebellar lob(ul)es, possible traces of their evolutionary history, and their genetic relationship with psychiatric disorders, cognitive performance and the cortical language network. We do so by leveraging publicly available GWAS summary statistics for imaging derived phenotypes (IDPs; based on the UK Biobank N*∼*31,000)[24], schizophrenia (SCZ)[25], autism spectrum disorder (ASD) [26] and cognitive performance[27] (Table S1).

## 2 Results

In order to gain further insights into possible genetic effects that go beyond the global cerebellar volume, we first performed an in-depth analysis of the genetic architecture of cerebellar substructures. A total of 32 cerebellar volumes were included in the analysis, including four global measures (left and right cerebellar cortical volume and cerebellar white matter, from the ‘aseg’ subcortical segmentation[28]), and 28 regional cerebellar measures (from the probabilistic cerebellar atlas [29]): left and right volume measures for lobules I-IV, V, VI, Crus I, Crus II, VIIB, VIIIA, VIIIB, IV, X, and vermis volume measures for VI, Crus I, Crus II, VIIB VIIIA, VIIIB, IX and X (see Figure S1).

In addition, we also used the data from [20], in which all cerebellar lobule volumes (from the probabilistic atlas [29]), with the exception of the vermis of Crus I, were included, to compute a total cerebellar volume measure and perform a GWAS for this global trait. These 33 cerebellar measures consisted of 19 independent variables, estimated with PhenoSpD [30] using a genetic correlation matrix of all cerebellar measures.

### 2.1 Cerebellar heritability and genetic correlations within the cerebellum

All cerebellar measures were heritable, with estimates ranging from 0.08 (se=0.015, p-val=1.4e-7) for the vermis of Crus I to 0.35 (se=0.015, p-val=1.4e-7) for the left and right cerebellar cortex (all corrected p-values *<* 0.05; see Table S2, Figure 1A). For each of the cerebellar substructures, we computed the genetic correlation between the vermis, left and right volumes, and tested whether they were significantly different to 1. Hemispheric volumes were highly correlated for all structures (r*_g_ >*0.9), indicating that most of the genetic effects are shared across cerebellar hemispheres. Nevertheless, the outermost lobules (X, IX and I-IV) had a genetic correlation estimate that was significantly lower than 1 (corrected p-values *<* 0.05, see Table S3), suggesting that there may be hemisphere-specific genetic effects on these substructures (Figure 1B). The genetic correlation between the vermis and the lateral cerebellar substructures was moderate (median r*_g_*=0.40, see Figure 1B), with the highest correlation for lobule IX (r*_g_*(Left-Vermis)=0.84, se=0.02, p-value=0), and the lowest for Crus I (r*_g_*(Right-Vermis)=0.11, se=0.08, p-value=0.13).

**Figure 1.**
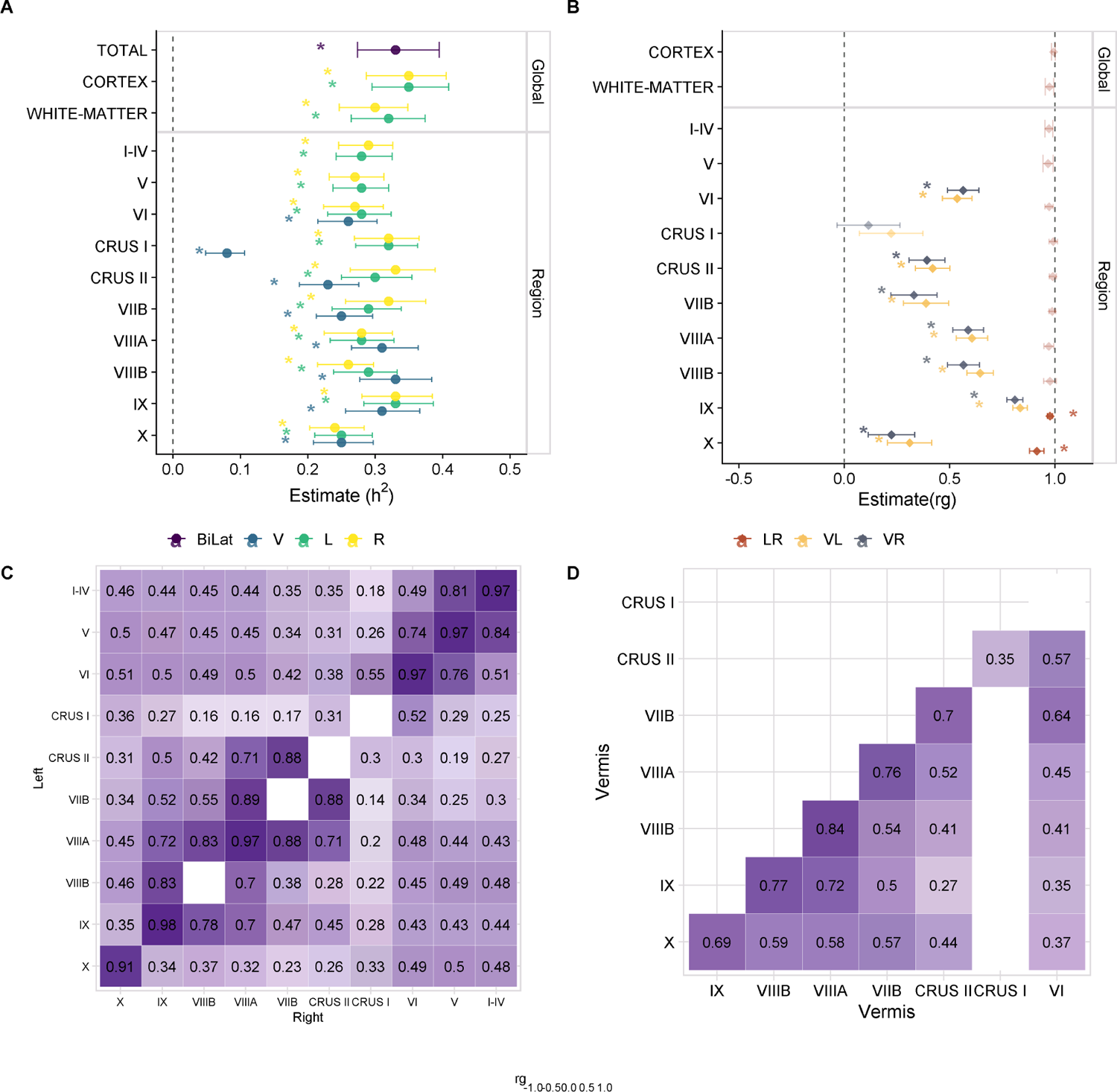
Genetic architecture of cerebellar substructures. **A** Heritability of cerebellar volumes. **B** Genetic correlations between volumes of a given cerebellar substructure. **C** Genetic correlations across cerebellar volumes within each hemisphere. Lower triangle shows the correlation pattern in the right hemisphere; upper triangle shows the correlation pattern in the left hemisphere; the diagonal shows the left-right genetic correlations. **D** Genetic correlations across cerebellar volumes in the vermis. For **A** and **B** estimates and 95% confidence intervals are shown. For **C** and **D** empty cells indicate non-significant estimates (after adjustments for multiple comparisons). h^2^= heritability; r*_g_*=genetic correlation; L= left; R= right; V= vermis; LR= left-right; VL= left-vermis; VR = right-vermis.

In order to assess whether there were distinct genetic effects on the cerebellum, we computed genetic correlations across cerebellar substructures within each hemisphere and within the vermis. This analysis revealed two main clusters that reflect cerebellar anatomy (Figure 1C): an anterior cluster, encompassing lobules I-IV, V, VI, Crus I and X, and a posterior cluster, consisting of lobules Crus II, VIIB, VIIIA, VIIIB and IX. Lobules within each cluster were highly correlated with each other (most r*_g_*s*>*0.5). Crus I was moderately correlated with both the anterior (r*_g_* range: 0.18-0.55) and the posterior (r*_g_*range: 0.14-0.36) clusters. The correlation pattern for the left and right hemispheres was very similar, while the pattern was a bit different for the vermis (Figure 1D): lobule X was genetically more correlated with the posterior cluster, while Crus I was only significantly correlated with Crus II (r*_g_*=0.4, se=0.09, adjusted p-value=0.0084).

### 2.2 Stratified heritability & evolutionary considerations

Next, we examined whether specific evolutionary genomic annotations (i.e. datasets consisting of genomic regions of evolutionary relevance) are depleted or enriched in heritability across the various cerebellar measures. We used stratified LDSC (S-LDSC) [31] to compute the contribution of variants within a given genomic annotation towards trait variation, and assess whether this contribution is larger or smaller than would be expected given the relative proportion of variants in that region. We considered six human-gained genetic and epigenetic sequence elements as genomic annotations marking different evolutionary periods (similar to the approach in [32]): adult and fetal Human Gained Enhancers[33], Human Accelerated Regions[34], Ancient Selective Sweeps[35], SNPs introgressed from other hominins [36] and genomic regions depleted of such introgression signals (so-called “introgression deserts”) [37]).

The vermis of Crus II showed a significant heritability depletion for so-called large introgression deserts (h^2^(C)=0.38, s.e.=0.15, adjusted p-value=0.0268, see Figure 2) [37]. No other cerebellar measure showed a significant enrichment or depletion in any of the annotations (Table S4).

**Figure 2.**
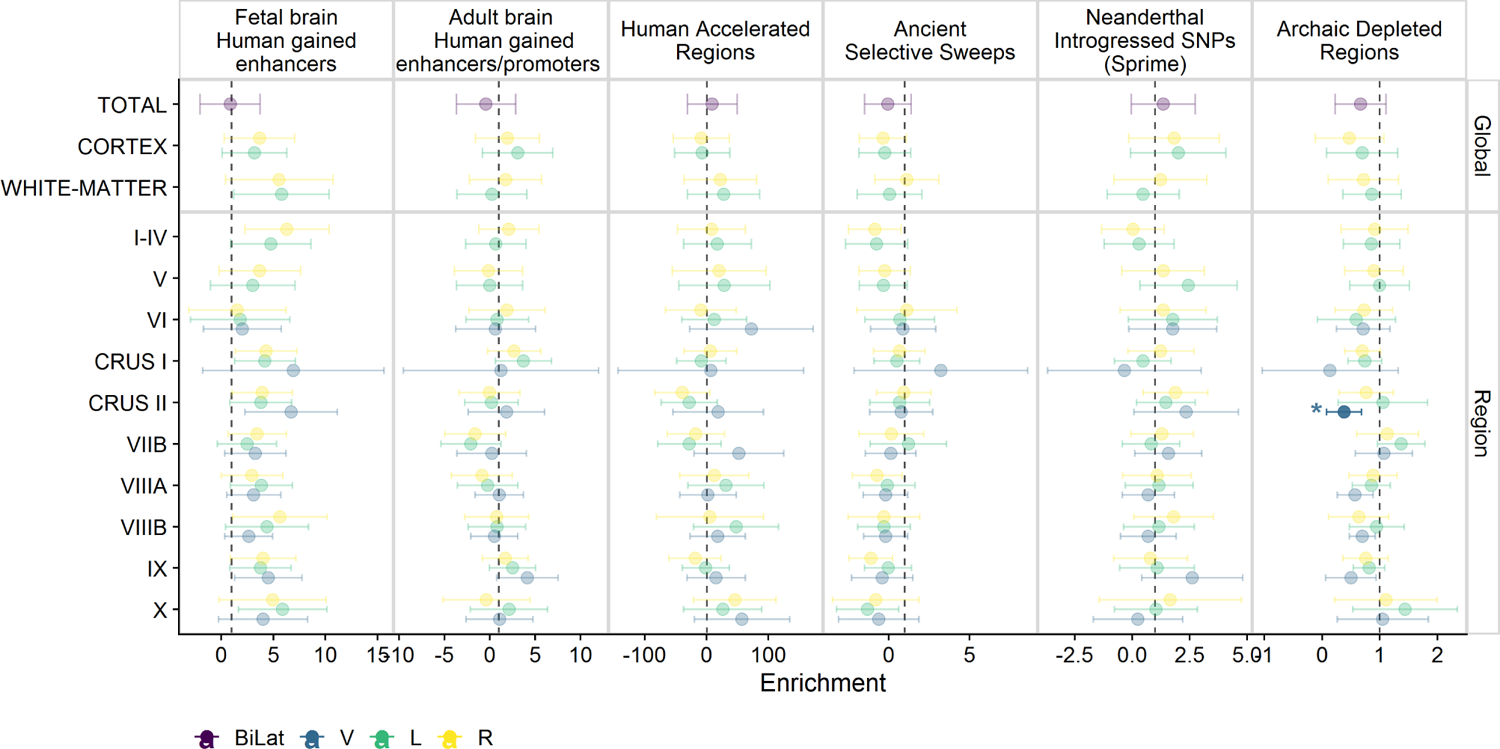
Stratified heritability analysis for six human-gained genetic and epigenetic sequence elements as genomic annotations marking different evolutionary periods. Each point reflects the enrichment estimate and error bars indicate the 95% confidence intervals. Estimates with a Bonferroni corrected p-value *<* 0.05 are highlighted with a brighter colour and marked with an *. Bilat=bilateral measure; V= Vermis; L= left; R=right.

### 2.3 Global genetic correlations with other brain regions, psychiatric disorders and cognitive traits

#### 2.3.1 Subcortical volumes

We found several genetic correlations between the cerebellar substructures and other subcortical volumes (‘Harvard-Oxford’ subcortical atlas) (Figure 3, Table S5). The brainstem, putamen and ventral striatum had positive correlations with most global cerebellar measures, except for cerebellar cortex volumes. Similarly, the regional cerebellar volumes had genetic correlations with the brainstem, putamen and ventral striatum, except for a few measures (i.e. Crus I and Crus II), and a lower genetic correlation with lateral VIIB and X lobules. We did not find any genetic correlations between the thalamus and global cerebellar volume, nor cerebellar cortex volumes, with estimates ranging from 0.04 to 0.10. There were, however, significant genetic correlations between the thalamic volumes and white-matter cerebellar volume (r*_g_* range: 0.26-0.31, all Bonferroni adjusted p-values<0.05). The pallidum was genetically correlated with the left and right cerebellar lobule I-IV, but not with the global measures. Overall, the genetic correlation patterns were quite stable across both hemispheres and the vermis, although there are also some noteworthy observations: bilateral lobule X measures were correlated with the brainstem but no other subcortical structures, while the vermis of lobule X was positively correlated with the putamen, caudate and ventral striatum.

**Figure 3.**
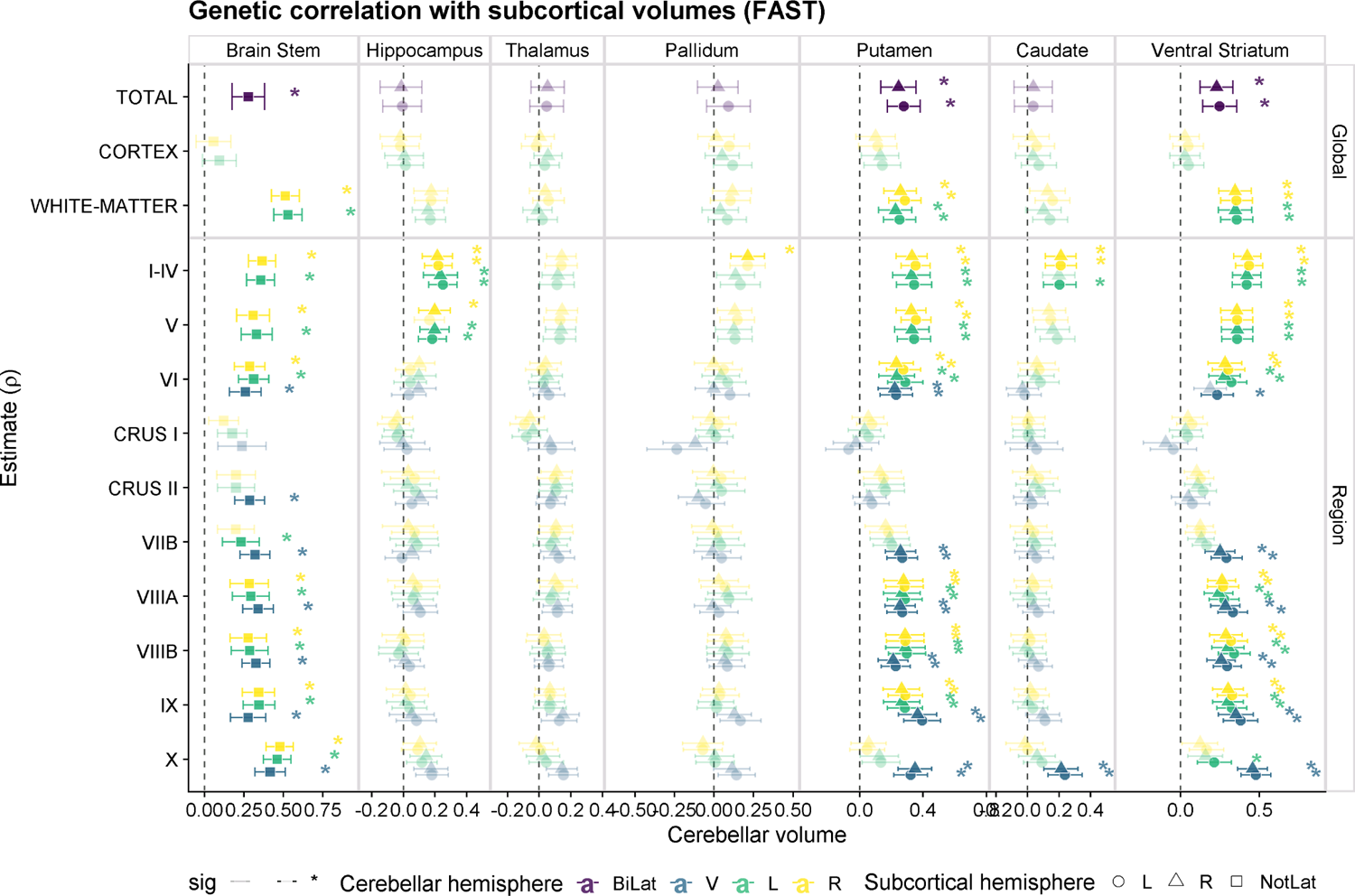
Genetic correlations between cerebellar measures and subcortical volumes (‘FAST‘ parcellation). Each point reflects the genetic correlation estimate and error bars indicate the 95% confidence intervals. Estimates with a Bonferroni corrected p-value *<* 0.05 are highlighted with a brighter colour and marked with an *. Bilat=bilateral; V= Vermis; L= left; R=right; NotLat= not lateralized.

To assess the robustness of these signals, and to enable a direct comparison with previous studies [20], we also computed the genetic correlations with subcortical volumes from the ‘aseg’ segmentation [28] (see Figure S3, Table S5): a similar trend can be observed for the brainstem, which also showed genetic correlations with the pallidum (most of the global measures except the cerebellar cortex and all the regional measures except for Crus I and Crus II) and the thalamus (cerebellar lobules I-IV, V and X)and less strong r*_g_*’s with the putamen or accumbens (as reported by [20] for the global cerebellar volume).

#### 2.3.2 Cortical language/reading network

We assessed genetic correlations between cerebellar volumes and cortical language and reading networks (Figure 4A). Total cerebellar volume had a positive r*_g_*with temporal occipital fusiform cortex in both hemispheres (left r*_g_*=0.27, se=0.06; right r*_g_*=0.22, se=0.06; adjusted p-values*<*0.05). Crus I and lobule VI volumes, but not Crus II, also showed significant positive global genetic correlations with regional volumes in the fusiform gyrus (i.e. temporal occipital fusiform and occipital fusiform gyrus) ipsi- and contra-laterally (r*_g_* range: 0.21-0.49, all corrected p-values*<*0.05). This association was present for both left and right cerebellar volumes but not with the vermis (Figure 4B). Left and right lobules I-IV were negatively correlated with the posterior part of the temporo-occipital fusiform cortex in both cortical hemispheres (r*_g_* range: −0.25; − 0.28). The vermis of IX had a r*_g_*of −0.30 (se=0.06; adjusted p-value*<*0.05) with posterior middle temporal gyrus (MTG); and lateral lobules VIIIB (but not the vermis) had a negative r*_g_* with the left temporo-occipital MTG (r*_g_*=0.35; r*_g_*=0.30). No other language-related cortical regions were genetically correlated with any of the cerebellar measures tested (Table S6).

**Figure 4.**
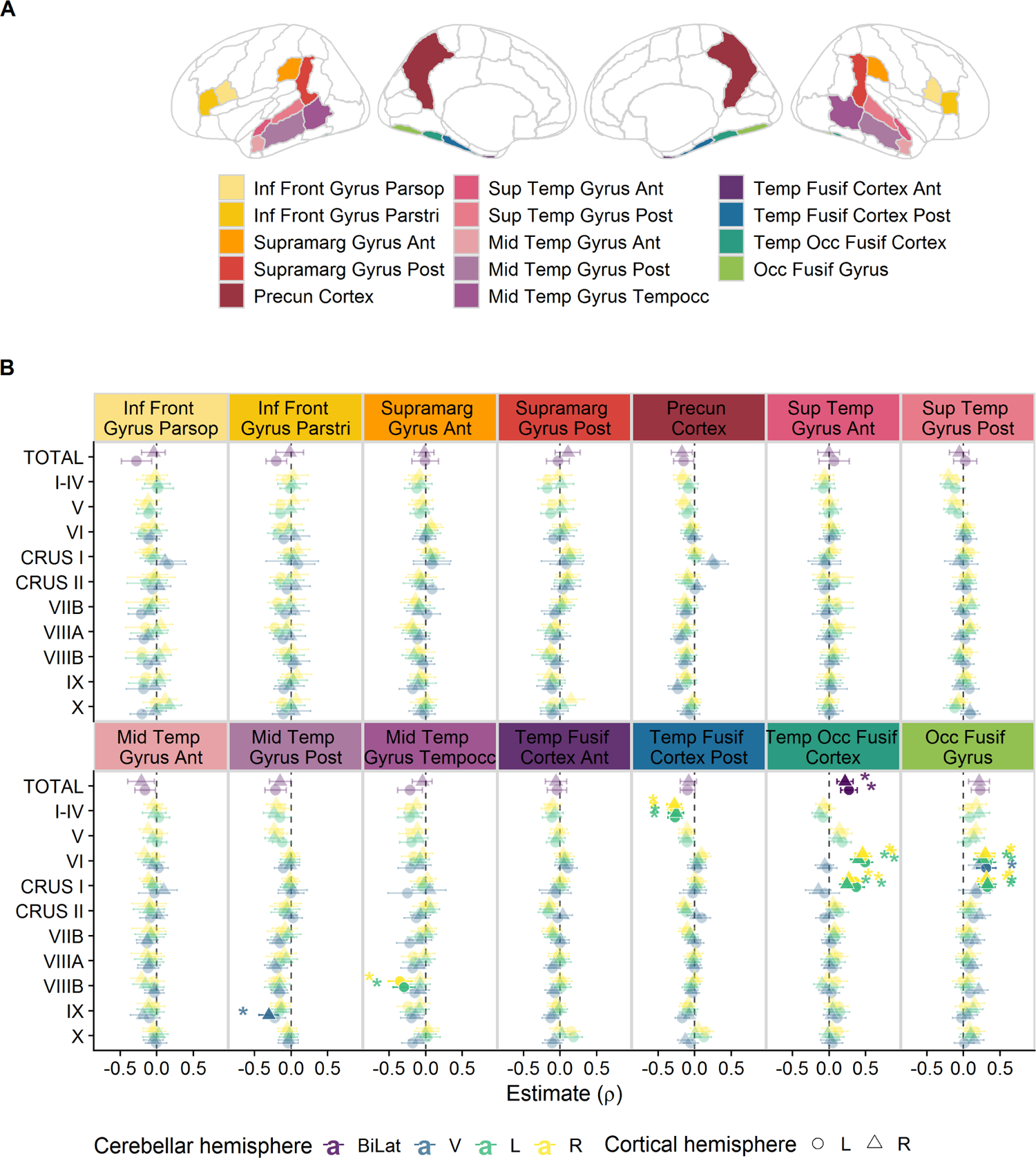
A. Cortical language network volumes used for the cortico-cerebellar genetic correlation analyses (regions from the Harvard-Oxford atlas). **B** Global genetic correlation between cerebellar volumes and cortical volumes. Estimates are depicted by points, and error bars indicate the 95% confidence interval. Estimates with a Bonferroni corrected p-value *<* 0.05 are highlighted with a brighter colour. Bilat=bilateral; V= Vermis; L= left; R=right.

#### 2.3.3 Psychiatric disorders and cognitive traits

Global genetic correlations of cerebellar lobules with ASD, SCZ and cognitive performance did not reveal any significant associations after multiple testing correction (all corrected p-values *>* 0.05) (Figure S2, Table S7).

### 2.4 Local genetic correlations

To probe some of the global genetic correlations further we computed local genetic correlations, which allow for positive and negative genetic correlations for different genomic loci, and are therefore more sensitive than global genetic correlations. Two different tools were used to compute local r*_g_*’s: LAVA[38] and SUPERGNOVA[39]. Local r*_g_*’s that were significant after multiple comparison correction (for both tools) are shown in Table 1, all local r*_g_*’s are shown in Table S8. We summarize the main results from LAVA here, and refer to SUPERGNOVA results in the Supplementary Material (Figures S4,S5, Table S8).

**Table 1.**
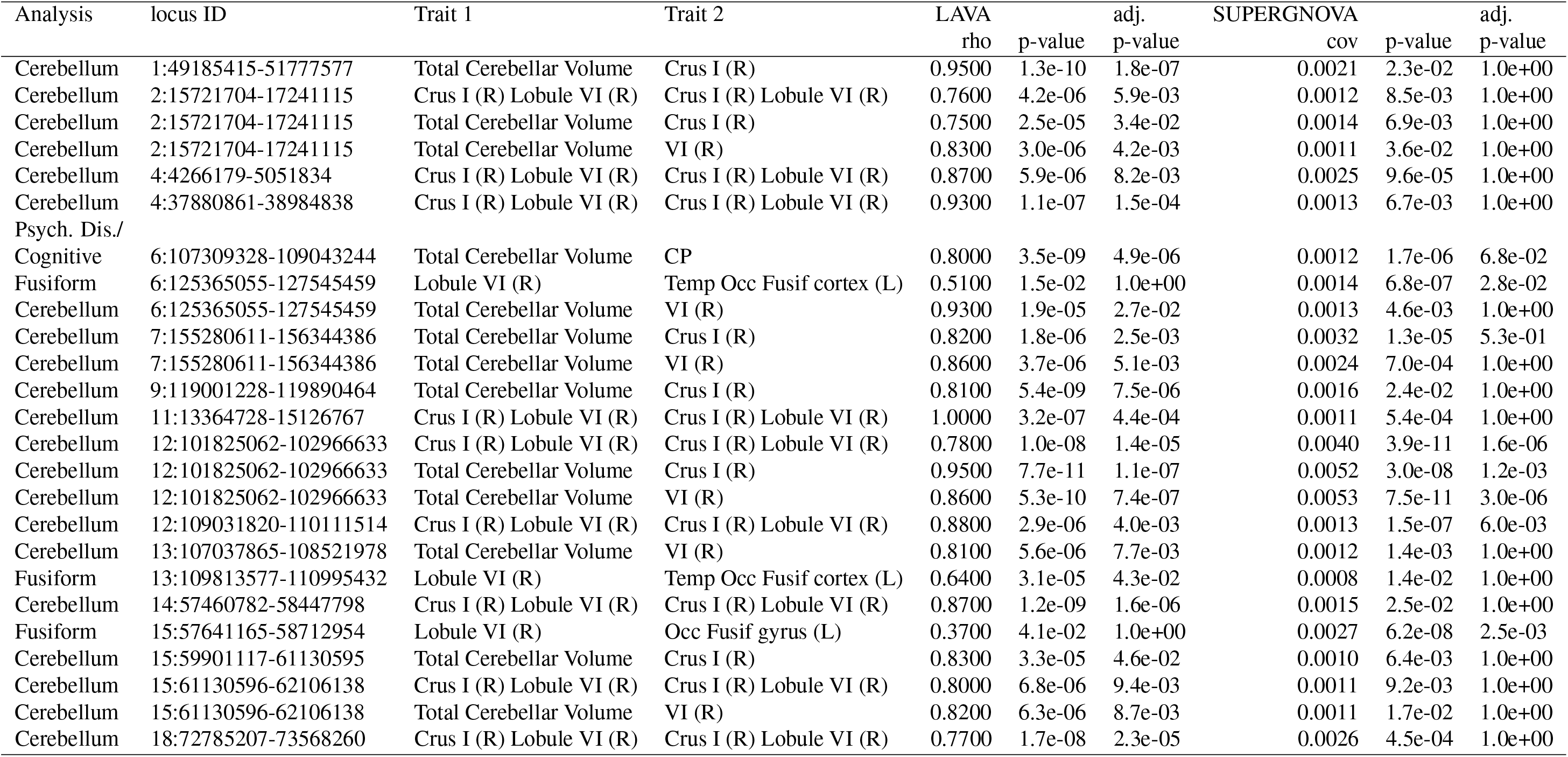
Local genetic correlations. Consistent signals across analysis tools (LAVA and SUPERGNOVA) are shown, i.e. estimates that were significant after Bonferroni multiple comparison correction in at least one of the tools and were nominally significant in the other one. LAVA: rho= local genetic correlation; p-value= p-value of local genetic correlation; adjusted p-value= p-value Bonferroni adjusted for 1,387 bivariate tests performed across 423 loci. SUPERGNOVA: cov= estimated local genetic covariance; p-value= The p value of local genetic covariance; adjusted p-value= p-value Bonferroni adjusted for 40,536 bivariate tests across 2,495 loci.

A three step approach was used to examine genetic correlations between three selected cerebellar measures and other traits. First, we computed the local genetic correlations between total cerebellar volume, right cerebellar Crus I and right lobule VI, which had 41 significant (adjusted p-values *<*0.05) local r*_g_*’s in 29 out of the 221 loci tested for these three trait-pairs using LAVA, with another 157 being nominally significant (Figure 5A and Table S8). All of these were positive correlations and contributed most to the global r*_g_* between these cerebellar traits (Figures S7-S35). Among these is a locus in chr9q33.1 (chr9:119001228-119890464), which included the top-hit of the total cerebellar volume GWASes[20, 21]. This locus includes the *PAPPA* gene, and showed local r*_g_* with Crus I but not lobule VI (Figure S18).

**Figure 5.**
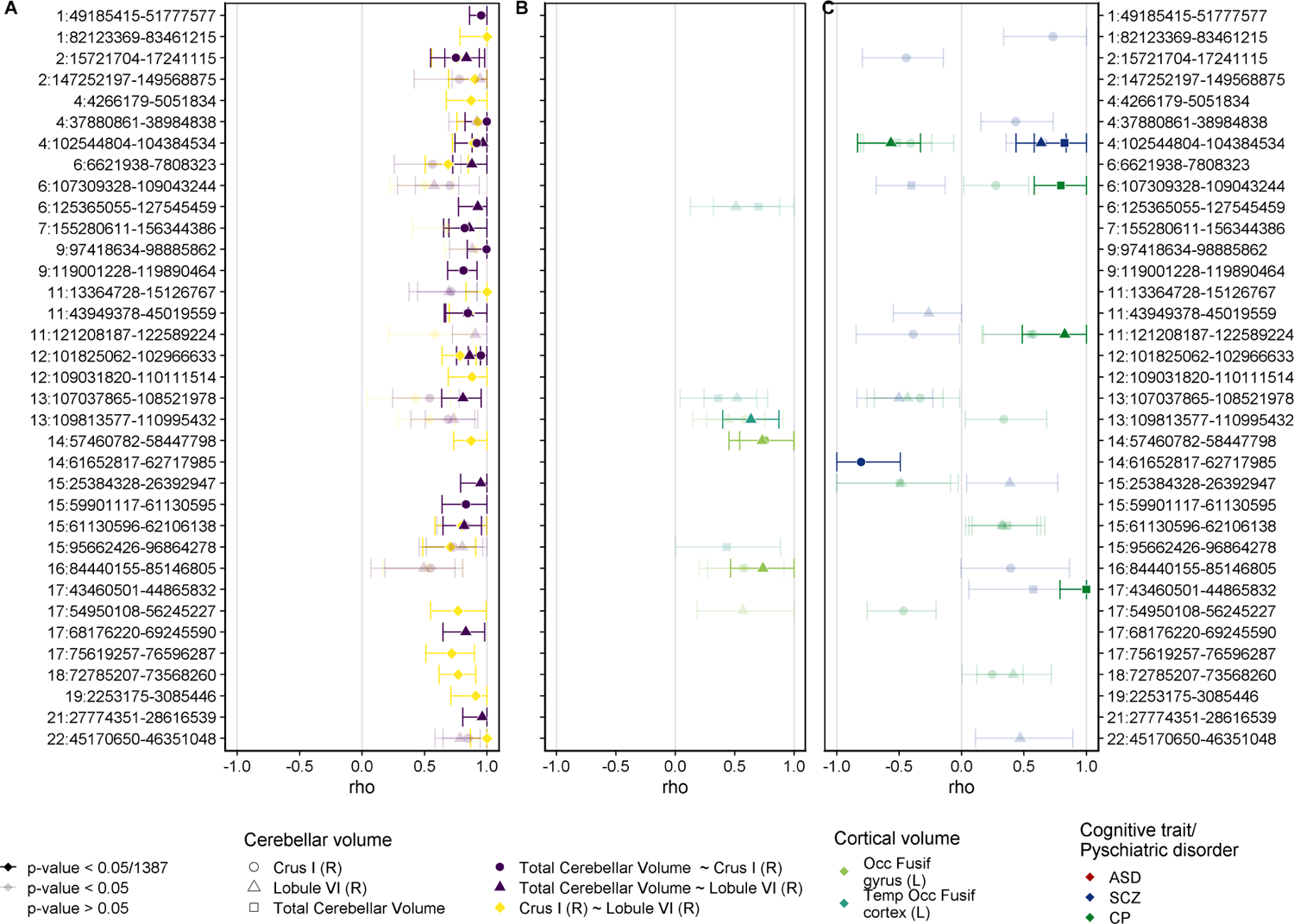
Local genetic correlations for genomic loci with significant non-zero estimates (using LAVA) after Bonferroni multiple testing correction. **A** local r*_g_* between cerebellar volumes: total, Crus I and lobule VI; local r*_g_* between these cerebellar volumes and **B** selected cortical volumes in the fusiform region or **C** cognitive traits/ psychiatric disorders. Each point reflects the estimate for a given pair of traits and locus. The error bars indicate the 95% confidence intervals. Bonferroni correction for multiple comparisons=0.05/1,387, as 1,387 local genetic correlations were run across all pairs (with non-zero univariate signal for both traits). rho= standardised coefficient for the local genetic correlation; R=right hemisphere; L=left hemisphere, Occ=occipital, Temp=temporal, Fusif=fusiform.

Next, we focused on the signal for global r*_g_* that was detected with two fusiform volumes (temporal occipital fusiform cortex, and occipital fusiform gyrus), and tested local r*_g_* between the total cerebellar volume, right Crus I and right lobule VI with the left cortical measures.This analysis revealed three loci showing significant local genetic correlations (13:109813577-110995432, 14:57460782-58447798, 16:84440155-85146805). A locus on chr14q22.3 had positive and significant genetic correlations with the occipital fusiform gyrus and both Crus I and lobule VI, but not with total cerebellar volume (Figures 5B,S24); the locus chr16:84440155-85146805, which was only significant for lobule VI and occipital fusiform gyrus (Figure S37); and the locus on chromosome 13q33.3-13q34, which had a significant r*_g_* between lobule VI and the temporal occipital fusiform cortex and a nominally significant r*_g_*between the occipital fusiform gyrus and right Crus I and lobule VI (Figure S36). These latter three associations were also nominally significant in the analysis using SUPERGNOVA (Table 1), and this locus on chromosome 13 was also positively genetically correlated with the three cerebellar measures tested using both methods, although it did not survive multiple comparison corrections for either.

Last, we assessed local correlations of these three cerebellar measures with ASD, SCZ and cognitive performance, to detect potential genetic correlations that may have been missed or cancelled out by the global genetic correlations (Figure 5C). There was no significant local r*_g_* between ASD and any of the three cerebellar measures tested, but there were multiple significant local r*_g_*’s with SCZ and cognitive performance in five different genomic loci (4q24, 6q21, 11q24.1, 17q21.31, 14q23.2, see Table S8). Multivariate analyses revealed that local r*_g_*s with SCZ and cognitive performance were mostly shared across the three cerebellar measures, reflected by diminished partial genetic correlations when adjusting for each other (Table S9). The strongest signal after adjustment was for the r*_g_* between total cerebellar volume and cognitive performance at the 6q21 locus (6:107309328-109043244: r*_g_*=0.80, adjusted p-value=4.87e-6), which was not affected when conditioning on lobule VI (partial correlation=0.8, p-value=0.000152), and slightly diminished when adjusting for Crus I (partial correlation=0.65, p-value=0.025). This bivariate signal was also captured by the SUPERGNOVA analysis, although it did not survive the multiple comparisons across the genome (uncorrected p-value=1.68e-6) (Table 1, Figure S38). The locus on chromosome 4q24 (4:102544804-104384534) showed an interesting pattern of genetic correlations: it had a positive local r*_g_* between the three cerebellar measures; a positive local r*_g_* between each cerebellar volume and schizophrenia, and a negative r*_g_* between the cerebellar measures and cognitive performance (nominally significant for Crus I and total cerebellar volume, significant after adjustments for lobule VI).

## 3 Discussion

Overall, our results confirm that the cerebellum is a heritable structure, both globally and at the regional level, and highlight the variability across its different substructures. They therefore support further investigation of the relationship between specific cerebellar substructures and other traits.

Within the cerebellum, global cerebellar measures were the most heritable ones. There was more variability for regional volumes, most of them were moderately heritable (h^2^ *>*0.2), except for the vermis of Crus I, which showed the lowest heritability, even if it was significantly different from zero. The vermis of Crus I, so called following the nomenclature that was used in the cerebellar probabilistic atlas [29], corresponds to the vermal component of VIIAf. This volume was excluded from previous studies that looked at total cerebellar volume due its small size (*<*0.005% of total cerebellar volume) [20, 40]. It is thus possible that the lower estimate for this region is driven by the larger measurement error associated with smaller volumes. Genetic correlations within the cerebellum reflected moderate to high genetic correlations across regions, which were very similar for both the left and right hemispheres and to some extent reflected the anatomical division of the anterior and posterior cerebellar lobes, which has been claimed to distinguish between the motor and affective/cognitive cerebellum [12]. Most of the cerebellar volumes had a perfect genetic correlation between the left and right volumes, while the vermal measures were all correlated with each other but only had a low to moderate genetic correlation with left and right hemispheric measures.

A depletion of heritability was detected for Crus II Vermis in the context of large regions or “deserts of introgression” [37]. These regions of the genome are depleted of introgressed archaic haplotypes and are enriched for genes expressed in the human brain, specifically in regions such as the cerebellum, the striatum and the thalamus [18, 41]. As has been suggested to explain the heritability depletion of cortical surface area measures in archaic deserts [42], the pattern observed here could be interpreted as the consequence of strong purifying selection against introgressed archaic variants, leading to a fixation of the modern variants in these “deserts of introgression” and hence a lower proportion of variance being explained by genetic variation within such regions. This explanation remains speculative but, in support of this idea, the vermis of Crus II also showed a positive enrichment of fetal brain human gained enhancers, although this did not survive multiple testing correction and multiple other cerebellar regions also show the same trend. The lack of enrichment of heritability in all other evolutionary annotations for all other cerebellar measures is in line with another study which, using an alternative approach, failed to find an enrichment of total cerebellar volume GWAS signal in human accelerated regions [21].

Global genetic correlations between total cerebellar volume and subcortical structures confirmed some of the previously reported genetic correlations[20], while our region-specific analysis provides greater resolution regarding the potential source of global cerebellar signals. There was a robust genetic correlation between cerebellar volumes and the brainstem, which was consistent across analyses (i.e. using different parcellations for the cerebellar structures), and replicated previous studies that used different GWAS summary statistics for the subcortical structures[20]. We did not find any significant genetic correlation between the thalamus and total cerebellar volume in either of the parcellations we used, in contrast to the previously reported genetic correlation of 0.24 between the thalamus (average left and right) and global cerebellar volume [20, 43]. There was, however, significant genetic correlation with some cerebellar lobules (I-IV, V and X). Further, we confirmed genetic correlations between the cerebellum and the pallidum[20], although there was variability with regard to the subcortical parcellation used for the pallidum and the specific cerebellar measures. This highlights that the genetic correlations that are currently being detected in genetic studies are sensitive to the specific datasets and/or the analysis parameters used. Analyzing specific lobules also revealed new genetic correlations between cerebellar measures and subcortical structures, including general positive genetic correlations between cerebellar lobules (except Crus I and Crus II) and the putamen and ventral striatum / accumbens; as well as some genetic correlations that were restricted to specific cerebellar lobes: such as lobules I-IV and V with the hippocampus, or lobules I-IV and vermal X with the caudate.

Cerebello-cortical genetic correlations, specifically targeting cortical volumes from the language and reading networks, revealed a strong genetic association between fusiform region volumes and cerebellar measures, total cerebellar volume and Crus I and lobule VI. This effect was not lateralized, as similar genetic correlations were observed both ipsi- and contralaterally, the strongest r*_g_* being between lobule VI and temporo-occipital fusiform cortex (r*_g_* range: 0.40-0.49, Bonferroni adjusted p-values*<*0.05). The vermis of lobule VI was also correlated with the left occipital fusiform gyrus (r*_g_*=0.31, adjusted p-value=1.48e-3). The fusiform cortex is known to be involved in visual processing, the left fusiform being involved in the orthographic mapping in reading, including word recognition. Interestingly, a meta-analysis reported reduced grey matter volume in bilateral lobules VI and right Crus II in dyslexia[44], while bilateral activation in lobule VI and Crus I has been linked to orthographic processing, with dyslexic children showing stronger functional connectivity between right cerebellar lobule VI and left fusiform gyrus during an orthographic task [45], although others report a lack of functional connectivity to the mid fusiform gyrus in the cerebellum [46].

It is important to stress that the nature of the present study is purely correlational, examining associations between genetic data and structural volume data. Thus, we cannot draw any direct links from the genetic correlation between cerebellar (lobule VI, Crus I) volumes and fusiform cortex to orthographic processing. Nevertheless, we hypothesize that shared genetic contributions to these specific cerebellar and cortical regions could impact their ability to optimally host processes in which both are involved, such as complex visual processing. Furthermore, local correlation analyses detected three possible relevant genomic loci that could contribute to this signal, two of which have previously been linked to dyslexia or language impairment: the locus in chr13q34 includes the gene *COL4A2*, which has been tentatively associated with dyslexia [47], and a second locus in 16q24.1 is a SLI linkage region and contains the gene *ATP2C2*, which has been associated with phonological short-term memory [48]. Of note, recent higher-powered studies have not replicated the impact of *COL4A2*, and have found only partial evidence for an association of *ATP2C2* with dyslexia but not with reading-related quantitative traits [49, 50].

We next assessed the global genetic correlations between the cerebellar measures and other traits, including ASD, SCZ and cognitive performance. In line with previous studies [20, 21], cerebellar volumes did not show whole-genome level correlation with ASD, SCZ or cognitive performance. Local genetic correlations revealed some evidence for pleiotropy: for instance, a locus in chromosome 6q21 was strongly correlated with cognitive performance and total cerebellar volume. This was the most robust signal, detected by both software programs used for the local genetic correlation analysis. Conditional analyses showed that this signal was dependent on genetic correlations with Crus I or lobule VI volumes, which were also nominally correlated with cognitive performance in this locus (Table S9). Regional association plots highlighted that the signal in this locus maps to the *FOXO3* gene. A query of the NHGRI-EBI GWAS Catalog[51] highlighted that multiple other traits including subcortical brain measures and psychiatric disorders were associated with this locus (ASD, SCZ, bipolar disorder and cross-psychiatric disorders). Furthermore, it has pleiotropic effects on SCZ and cognitive abilities[52]. In line with this, our local genetic correlation analysis also pointed to a negative genetic correlation between total cerebellar volume and SCZ in this locus, although this signal did not survive multiple testing correction.

A further locus in chromosome 4q24 showed positive correlations with total cerebellar volume, Crus I and lobule VI; and also between each of these cerebellar volumes and schizophrenia, while being negatively correlated with cognitive performance. This locus is a genome-wide significant locus for all traits except Crus I (R), and it has been associated with multiple other traits, including subcortical volumes and cerebellar measures[24], and psychiatric disorders. Conditional analyses confirmed that the genetic correlations with SCZ and cognitive performance capture shared signals between the three tested cerebellar measures.

Three other local genetic correlations were significant for one pair of cerebellar measures and SCZ or cognitive performance, each of these local r*_g_*’s being independent of the other cerebellar measures we tested in the conditional analyses. The negative genetic correlation between Crus I and SCZ in 14q23.2 seems potentially specific: this locus encompasses the *SYT1* gene, which encodes for Synaptotagmin I, an integral membrane protein of synaptic vesicles that is involved in calcium-dependent neurotransmitter release [53]. Although *SYT1* has not been directly implicated in SCZ, it is downstream of the SCZ risk factor *MIR137*[54]. The locus in chr17q21.31 corresponds to the common inversion region of high linkage disequilibrium including *MAPT* and *KANSL1* and is also associated with many traits, including neurodegenerative diseases such as Alzheimer’s and Parkinson’s diseases[55, 56], and behavioural traits such as intelligence and handedness[57].

In sum, despite the lack of global genetic correlation, local genetic correlations enabled us to pinpoint some interesting genomic loci that may have pleiotropic effects between cerebellar measures and SCZ or cognitive performance.

Two caveats should be noted regarding the present study. First, it relied heavily on the UK Biobank dataset for all the neuroimaging traits analyzed. Although this is a powerful resource and one of the largest and most homogeneous brain imaging genetics datasets to date [24], participation bias in the UK Biobank is known to distort the genetic correlation estimates [58, 59]. Furthermore, we leveraged summary statistics that were readily available from previous studies[24], which restricted our ability to assess the sensitivity of the results (e.g. by including or excluding covariates to identify potential confounds or collider effects). The results presented in this study should therefore be replicated in studies using different designs that would enable their generalizabilty to be tested. Second, we focused on volumetric measures of cerebellar lobules which do not fully reflect functional regions [7]. A task-based functional parcellation of the cerebellum recently provided a detailed mapping of its functional organization, and highlighted its involvement in cognition [7]. Using these functionally defined regions in future studies will therefore provide additional insights into the genetic relationship between different cerebellar regions and other brain, psychiatric or cognitive phenotypes.

In summary, we have performed a comprehensive genetic analysis of cerebellar volumes, which has provided new insights regarding the genetic relationships that are shared or unique between specific lobules and other brain regions and traits. Interestingly, our results highlighted lobule VI, Crus I and Crus II. These cerebellar regions have been consistently linked to cognitive functions such as language, and with social cognition in general (mentalizing, social action sequencing), as well as showing a general involvement in executive control and attention [60, 61, 62, 63]. It is also noteworthy that our study failed to detect strong lateralization effects, possibly pointing to the relevance of both hemispheres even in domains (such as speech) where lateralization effects are expected [64], although this could also indicate a lack of sufficient resolution when relying on structural data.

## 4 Methods

### 4.1 Data

GWAS summary statistics for imaging derived phenotypes (IDPs) based on the UK Biobank (N*∼* 31,000) were downloaded from the Oxford Brain Imaging Genetics Server – BIG40 (https://open.win.ox.ac.uk/ukbiobank/big40/) [24]. These IDPs included a total of 33 cerebellar measures: 10 volumes of cerebellar regions (28 measures in total: left, right and vermis for 8 measures, left and right only for 2 measures; Harvard-Oxford subcortical parcellation, plus two global measures: cerebellar cortical volume and cerebellar white-matter volume (left and right) [28]. In addition, we also included the summary statistics from the recently published total cerebellar volume GWAS, in which total cerebellar volume had been computed as the sum of all the aforementioned FAST cerebellar volumes except for the Crus I vermis volume [20]. Additional brain volumes were also used for some analyses (see Table S1 for all measures). These included subcortical volumes (13 subcortical volumes from the Harvard-Oxford atlas; 15 from the subcortical atlas ‘aseg’[28]), and the 28 cortical volumes (Harvard-Oxford atlas [65]) of regions of interest related to language (14 left and 14 right volumes; see Figure 4A).

All GWAS summary statistics used in the current study are specified in Table S1. For each GWAS summary statistic dataset, we kept unique unambiguous SNPs and merged these with the HapMap3 SNPs. All datasets adopted hg19 genomic coordinates.

### 4.2 Estimating number of independent cerebellar traits

PhenoSpD was used to define the number of independent cerebellar traits across these 33 measures[30, 66]. PhenoSpD uses GWAS summary statistics to first estimate the phenotypic correlations across traits, and then applies spectral decomposition of matrices to identify the number of independent variables [30, 66]. We used the estimated effective number of independent variables (‘VeffLi’) to adjust using Bonferroni correction for multiple comparisons across the independent cerebellar measures (see Figure S1).

### 4.3 SNP-based heritability

SNP heritability is the proportion of variance explained by common genetic factors, and was computed using GWAS summary statistics by running the LDSC (v1.0.1) tool [67]. The significance of the heritability estimates was Bonferroni corrected for multiple comparisons for the 19 independent cerebellar measures as defined by PhenoSpD (see above; p-value thresh-old=0.05/19=0.0026).

### 4.4 Stratified heritability analysis

We used stratified LDSC (S-LDSC) [31] to compute the contribution of variants within each specific genomic region towards trait variation, and assess whether this contribution was larger or smaller than would be expected given the relative proportion of variants in that region. We considered six human-gained genetic and epigenetic sequence elements as genomic annotations marking different evolutionary periods (similar to the approach taken in [32, 68]): fetal brain Human Gained Enhancers (HGE) [33], adult HGE and promoters in the cerebellum [69], Human Accelerated Regions [34], Ancient Selective Sweeps [35], SNPs introgressed from other hominins [36] and genomic regions depleted from such introgression signals (so-called “introgression deserts”) [37]).

For each of these evolutionary categories, annotations and LD-scores were created following instructions from the LDSC wiki (https://github.com/bulik/ldsc/wiki/LD-Score-Estimation-Tutorial). We then estimated heritability enrichment or depletion for each category, using the baselineLDv2.2 model (which includes 97 annotations including several other regulatory elements, linkage statistics and measures of selective constraint). For human gained enhancers and promoters (i.e. fetal HGE and adult human gained enhancers and promoters), epigenetic marks from the fetal brain (E081 and E082) and adult brain (E073) from the Epigenome Roadmap Project 25 state model were also included in the model [70].

### 4.5 Global genetic correlations

The genetic correlation (r*_g_*) between two traits is the proportion of shared variance explained by common genetic factors. LDSC (v1.0.1) was used to estimate global genetic correlations between pairs of traits using GWAS summary statistics [67]. For every pair of measures for the left and right volumes of each cerebellar structure, we tested whether r*_g_* was significantly different to 1, whereas for analyses involving different cerebellar measures or non-cerebellar measures, we tested whether the genetic correlation estimate was significantly higher than 0.

Genetic correlations within the cerebellum were computed for a given cerebellar volume between left-right, left-vermis and right-vermis volumes; for the different volumes within the left-hemisphere, right-hemisphere and vermis (200 r*_g_* in total, p-value threshold=0.05/200=0.00025).

Cerebellar volumes were also tested for genetic correlations with subcortical measures, cortical volumes within the language network, ASD, SCZ and cognitive performance. Multiple comparison corrections were applied using the Bonferroni method: correcting for 19 independent cerebellar traits and the number of other traits within each analysis, which were: 3 cognitive/disorder traits(p-value threshold=0.05/(19*×*3)=0.00088); 13 (‘Harvard-Oxford’) or 15 (‘aseg’) subcortical volumes (p-value threshold=0.05/(19*×*13)=0.0002; p-value threshold=0.05/(19*×*15)=0.00017); 28 cortical volumes (p-value threshold= 0.05/(19*×*28)=9.4e-05).

### 4.6 Local genetic correlations

The global genetic correlation is an average correlation of genetic effects across the genome, and can therefore cancel out local genetic correlations with opposite direction of effects across different genomic loci. The local genetic correlation partitions the genome into *∼* 2500 similar-sized chunks and computes the genetic correlation between two traits separately for each of these genomic loci, allowing for local genetic correlations with opposite directions across loci between two traits.

We used LAVA [38] to compute local genetic correlations between total cerebellar volume and right-cerebellar (Crus I and lobule VI) and left-cortical volumes (14 language- and reading-related regions in the left hemisphere, Figure 4A), using a three-step analysis. First, local univariate analysis was run for each trait to identify loci with univariate genetic signal (p*<* 1 *×* 10*^−^*^4^). Next, we ran local genetic correlations for each pair of traits in the loci that had non-zero univariate estimates for both traits. A total of 1,387 bivariate tests were performed across 423 loci S8. This approach allowed us to identify which genetic loci were contributing most to the global r*_g_*. The regional associations for each of the traits that contributed to significant local genetic correlations were visualized using ‘gassocplot’ package in R (v 4.1.1) [71, 72]. Last, each of the seven significant local genetic correlation signals between a cerebellar trait and psychiatric disorder or cognitive traits were conditioned on the other two cerebellar measures (if univariate analysis was significant for the conditioning phenotype) to obtain partial genetic correlations.

For sensitivity, we ran the same pairwise bivariate local genetic correlations using SUPERGNOVA [39], which is another tool for computing bivariate local genetic correlations, but does not perform multivariate analysis. The same genome partition definition (i.e. file ‘blocks_s2500_m25_f1_w200.GRCh37_hg19.locfile’ as provided by LAVA) was used for SUPERGNOVA to facilitate comparison of the identified genomic loci (Figure S6). A total of 40,536 local genetic correlations were run across all pairs of traits and 2,495 genomic loci using SUPERGNOVA. We then assessed whether the two methods provided concordant results, to identify the most robust signals.

## Data availability

GWAS summary statistics used in this study (listed in Table S1) are available from the NHGRI-EBI GWAS Catalog and from Oxford Brain Imaging Genetics Server - BIG40.

## Code availability

The custom code associated with this study is publicly available at https://github.com/amaiacc/MS-cerebellum-UKB.

## Acknowledgements

A. C-C. is cofunded by the Spanish Ministry of Science and Innovation and the Agencia Estatal de Investigación through Ayudas Juan de la Cierva-Incorporación (ref. IJC2018-036023-I) and supported by the Programa Fellows Gipuzkoa de atracción y retención de talento from the Diputación Foral de Gipuzkoa. BCBL acknowledges funding from the Basque Government through the BERC 2022-2025 program and by the Spanish State Research Agency through BCBL Severo Ochoa excellence accreditation CEX2020-001010-S. CB acknowledges support from the Spanish Ministry of Science and Innovation (grant PID2019-107042GB-I00), MEXT/JSPS Grant-in-Aid for Scientific Research on Innovative Areas #4903 (Evolinguistics: JP17H06379), and the support of a 2020 Leonardo Grant for Researchers and Cultural Creators, BBVA Foundation. Funding bodies take no responsibility for the opinions, statements and contents of this project, which are entirely the responsibility of its authors.

## Supplementary information

### Supplementary tables

**Table S1.** Publicly available genome wide association study (GWAS) summary statistics used in the current study.

**Table S2.** Heritability estimates for 33 cerebellar measures.

**Table S3.** Genetic correlation estimates within the cerebellum.

**Table S4.** Heritability enrichment results for evolutionary annotations.

**Table S5.** Genetic correlation estimates between cerebellar and subcortical volumes.

**Table S6.** Genetic correlation estimates between cerebellar and language network cortical volumes.

**Table S7.** Genetic correlation estimates between cerebellar volumes and psychiatric disorders and cognitive traits.

**Table S8.** Local genetic correlation estimates across genomic loci (LAVA and SUPERGNOVA).

**Table S9.** Conditional local genetic correlation estimates (LAVA).

### Supplementary figures

**Figure S1.**
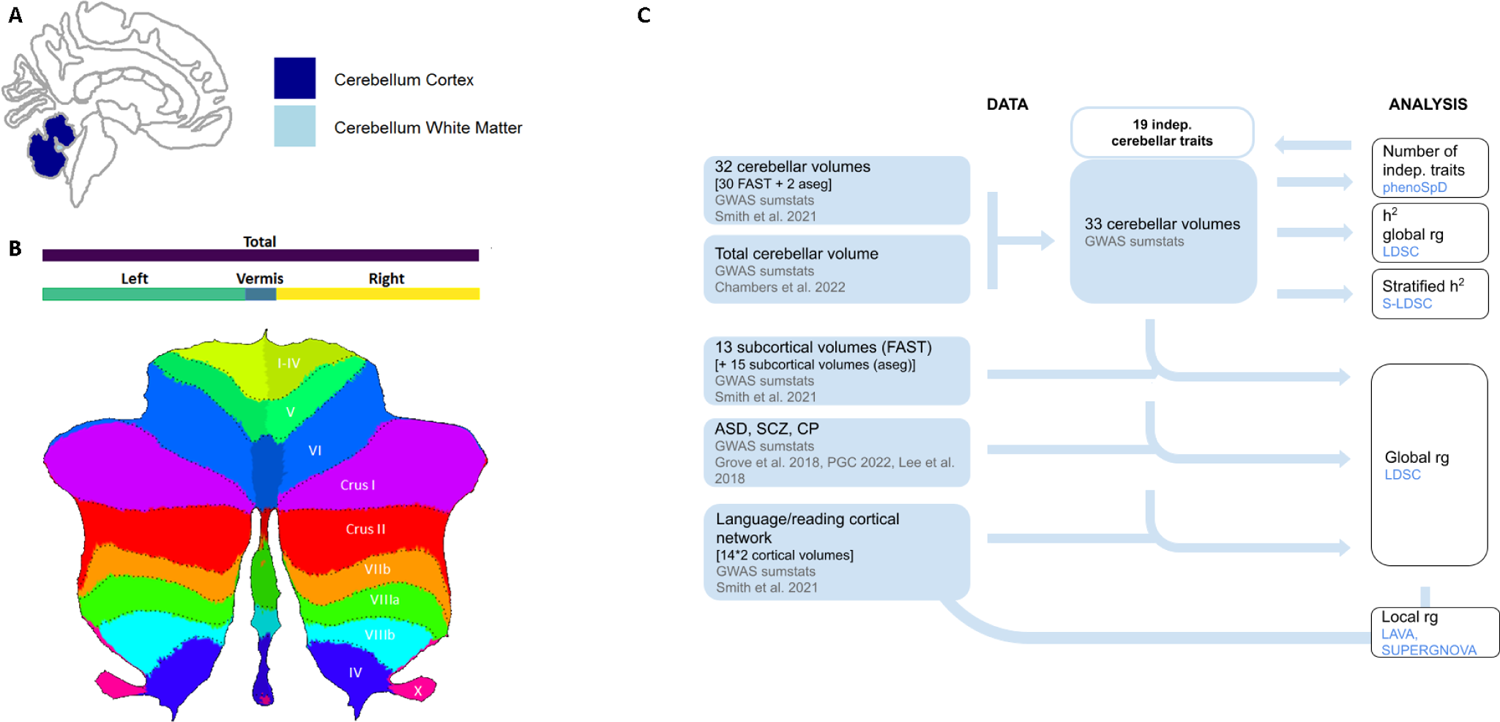
Cerebellar measures included in the study: **A** cerebellar cortex and white matter (‘aseg’ atlas, left and right), **B** total cerebellar volume4 and cerebellar lobules (‘Diedrichsen atlas SUIT’[29], left, right and vermis) and **C** study workflow including datasets and analytical approach. GWAS sumstats=Genome Wide Association Study summary statistics, phenoSpD=phenotypic spectral decomposition; h^2^=SNP-heritability; rg= genetic correlation; LDSC= linkage disequilibrium score regression; S-LDSC= stratified LDSC.

**Figure S2.**
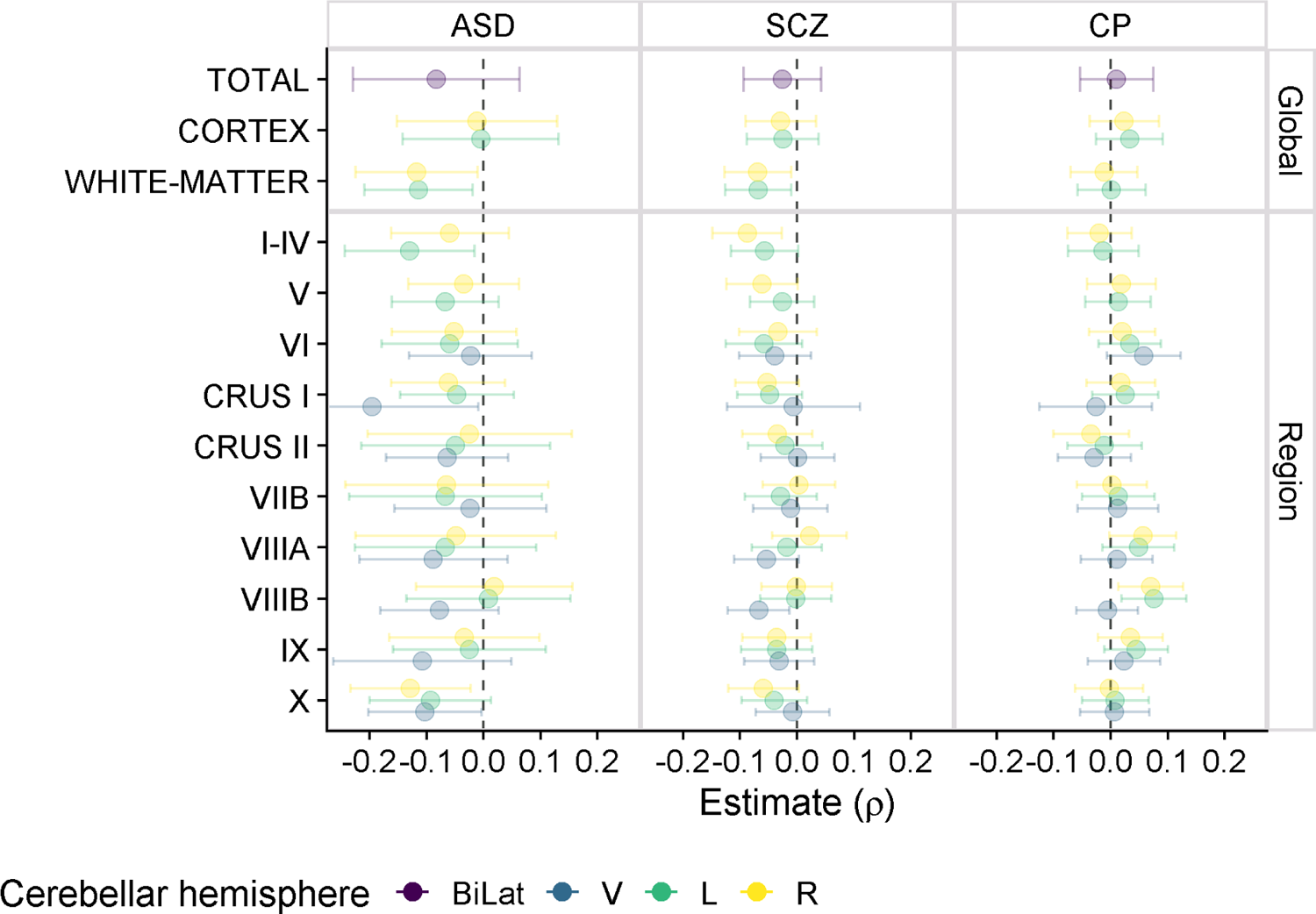
Global genetic correlation with psychiatric disorders and cognitive traits. Estimates are depicted by points, and error bars indicate the 95% confidence interval. All estimates had a Bonferroni corrected p-value *>* 0.05. ASD=Autism Spectrum Disorder; SCZ=Schizophrenia; CP=cognitive performance. Bilat=bilateral measure; V= Vermis; L= left; R=right.

**Figure S3.**
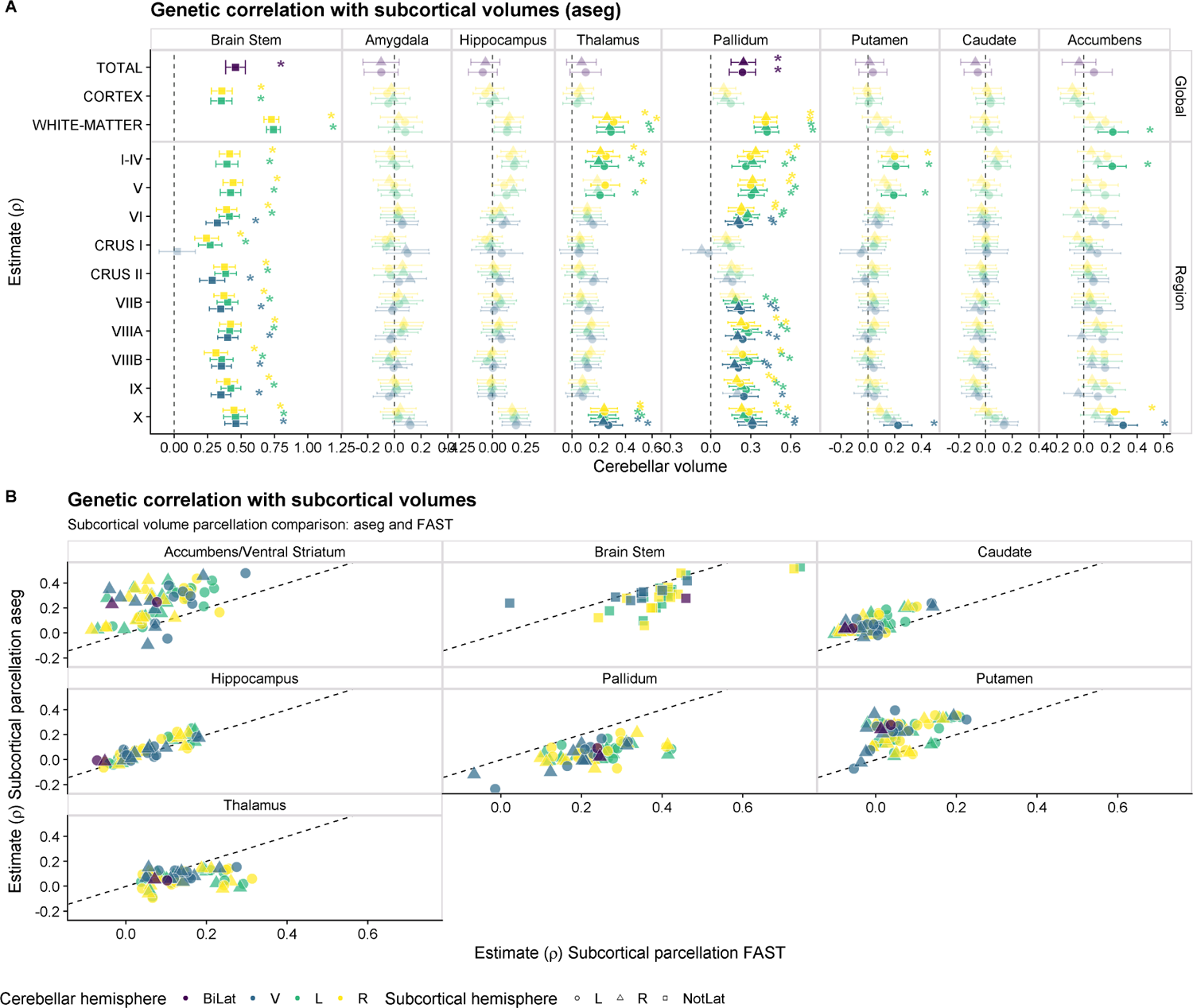
Genetic correlation with subcortical volumes. **A** Genetic correlation between cerebellar measures with subcortical volumes derived using ‘aseg’ parcellation. B Comparison of the genetic correlation between cerebellar volumes and the two subcortical parcellations (‘FAST‘ and ‘aseg‘) for each subcortical structure.

**Figure S4.**
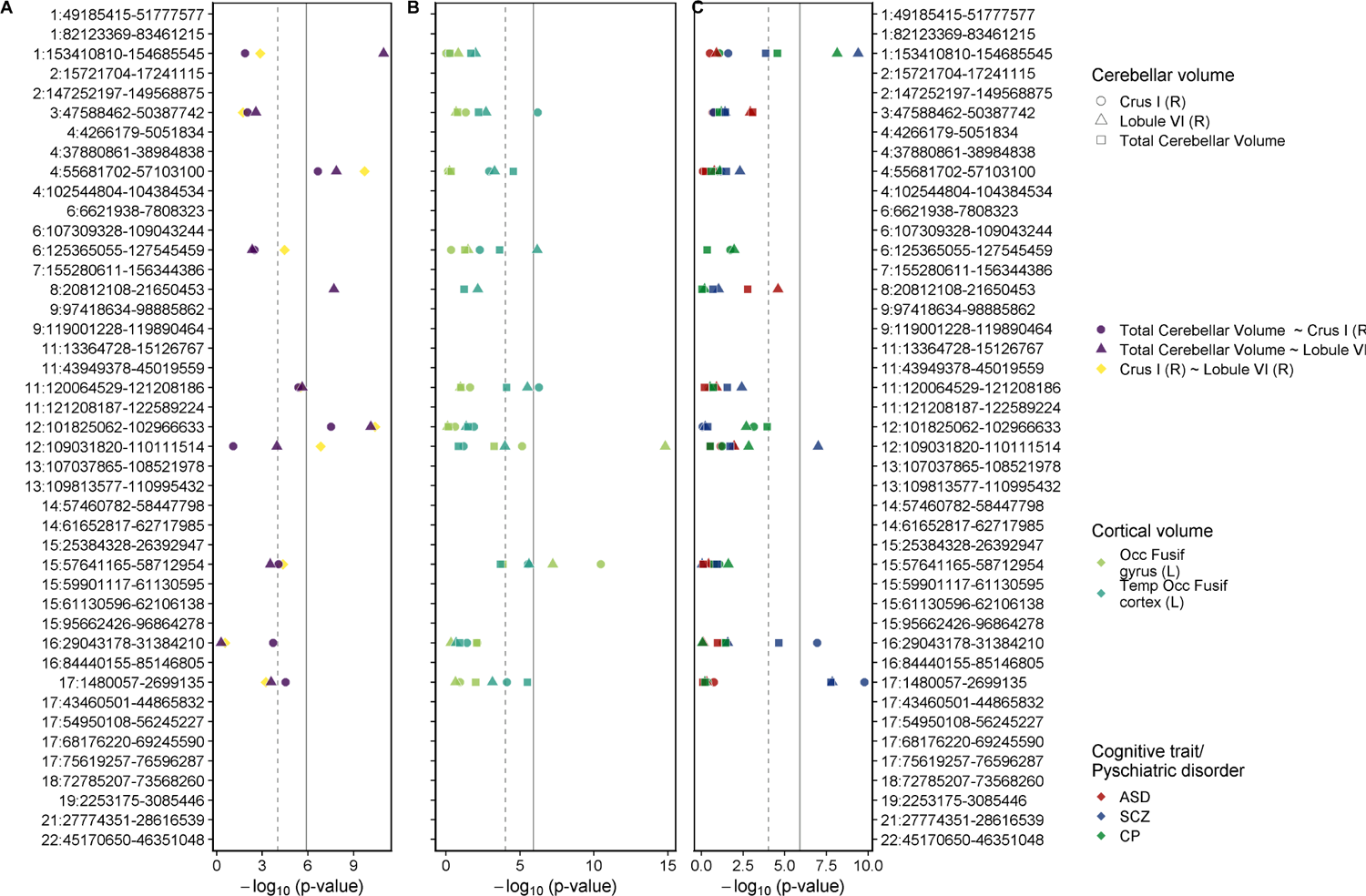
P-values associated with the local genetic covariances (using SUPERGNOVA) across the genome between **A** cerebellar volumes Crus I and lobule VI, and between each of these two cerebellar measures with **B** selected cortical volumes in the fusiform region and **C** cognitive traits/ psychiatric disorders. Only local genetic correlations with non-zero heritability estimates per trait are shown. Bonferroni correction for multiple comparisons=0.05/40,536, as 40,536 local genetic correlations were run across all pairs and 2,495 genomic loci. R=right hemisphere; L=left hemisphere, Occ=occipital, Temp=temporal, Fusif=fusiform.

**Figure S5.**
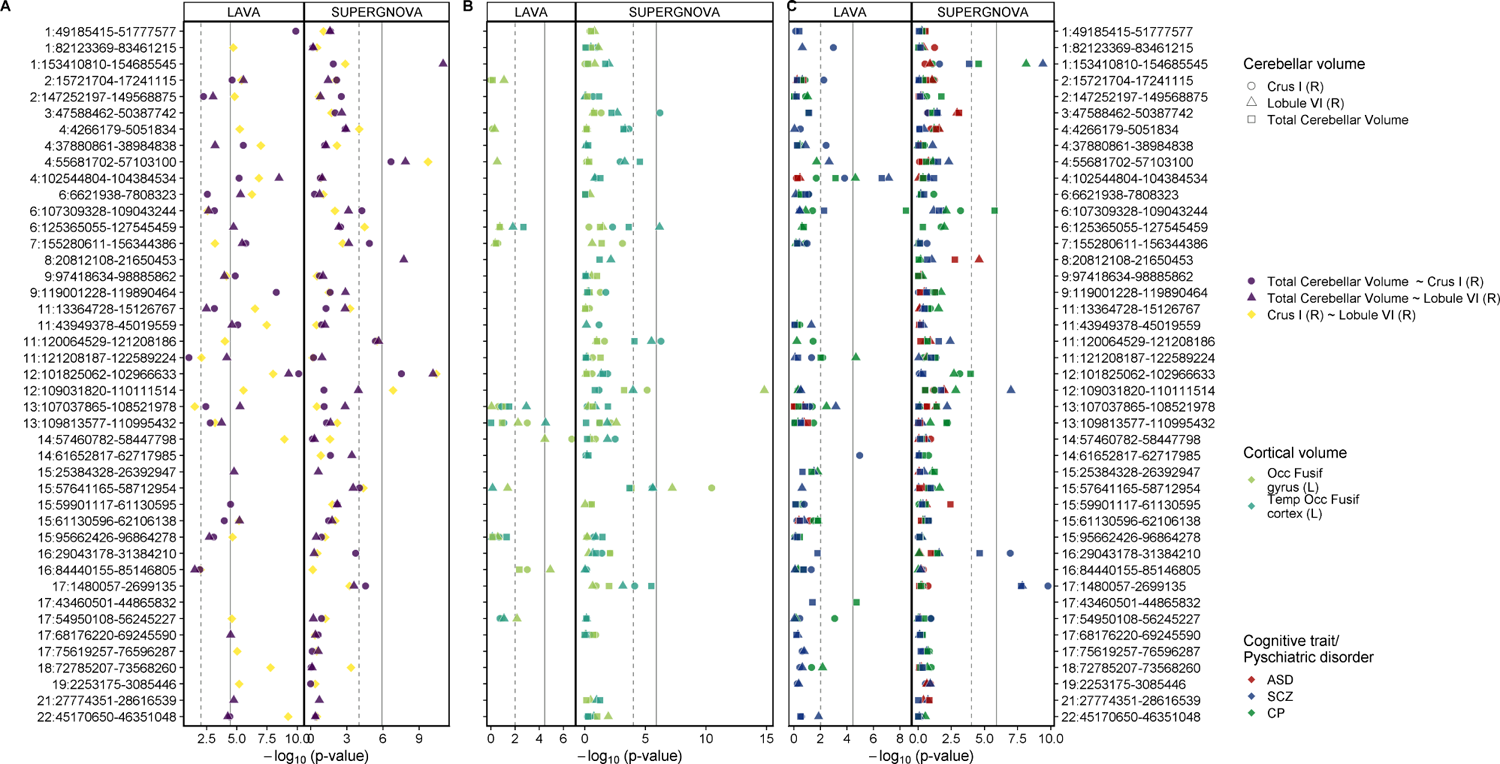
Local genetic correlations (using LAVA and SUPERGNOVA) across the genome between **A** cerebellar volumes Crus I and lobule VI, and between each of these two cerebellar measures with **B** selected cortical volumes in the fusiform region and **C** cognitive traits/ psychiatric disorders. Only genetic loci that showed a significant local rg in one of the tests are shown. The dashed lines indicate the p-value=0.05. The solid grey line indicates Bonferroni correction threshold for multiple comparisons in each method (1,387 tests in LAVA, 40,536 tests in SUPERGNOVA). R=right hemisphere; L=left hemisphere, Occ=occipital, Temp=temporal, Fusif=fusiform.

**Figure S6.**
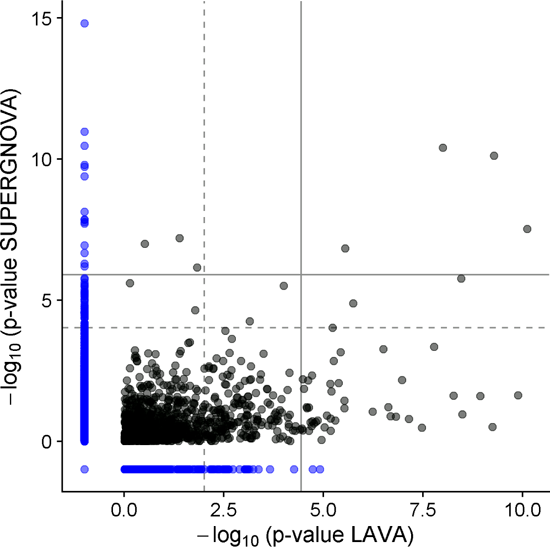
Local genetic correlation p-value comparison across tools: LAVA and SUPERGNOVA for the same genomic partition. The solid grey line indicates Bonferroni correction threshold for multiple comparisons in each method (1,387 tests in LAVA, 40,536 tests in SUPERGNOVA). The blue dots indicate loci that were only considered in one of the tools.

**Figure S7.**
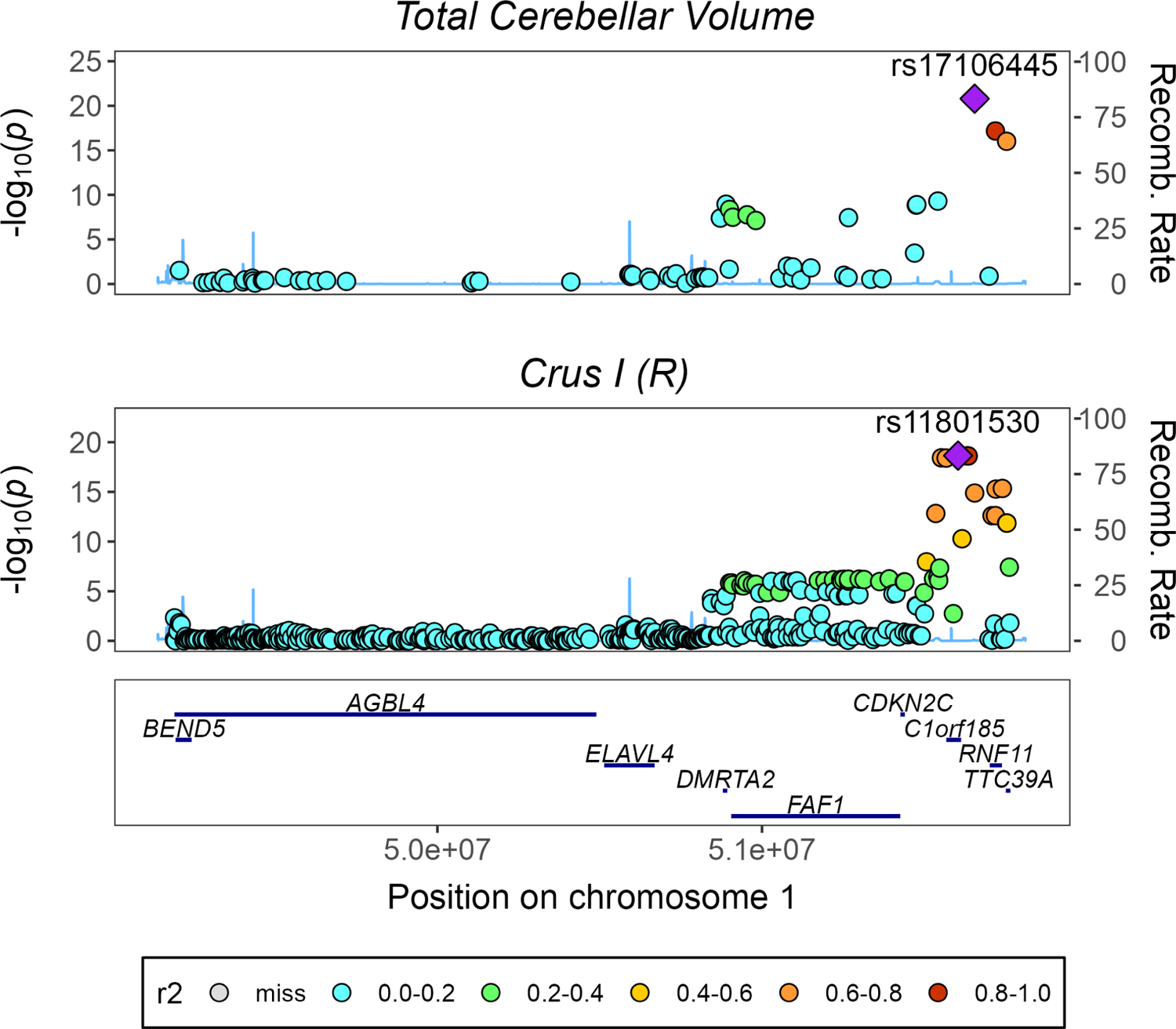
Regional association plot for traits with a significant local genomic correlation in locus 1:49185415-51777577, (LAVA). HapMap3 SNPs are shown.

**Figure S8.**
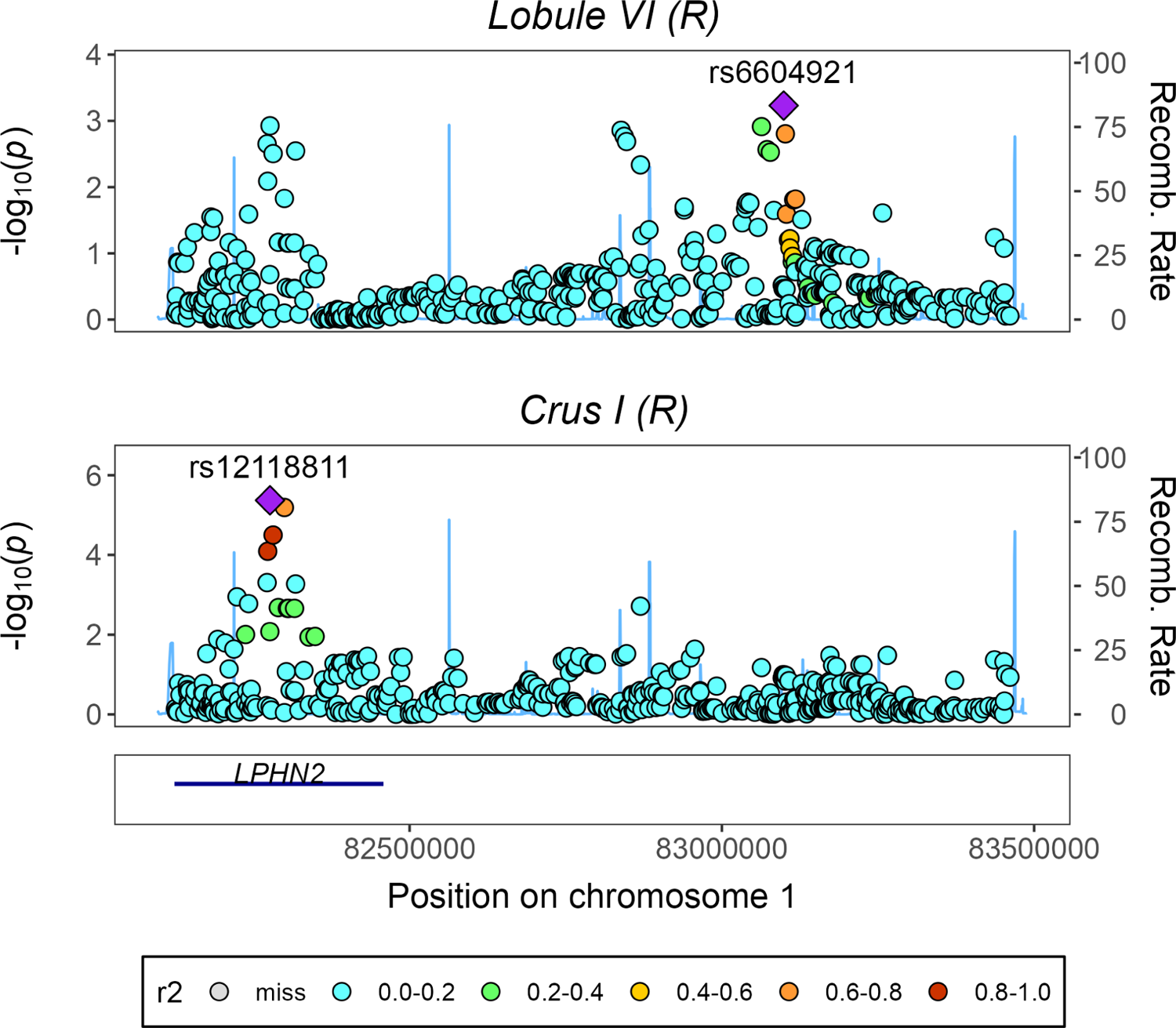
Regional association plot for traits with a significant local genomic correlation in locus 1:82123369-83461215, (LAVA). HapMap3 SNPs are shown.

**Figure S9.**
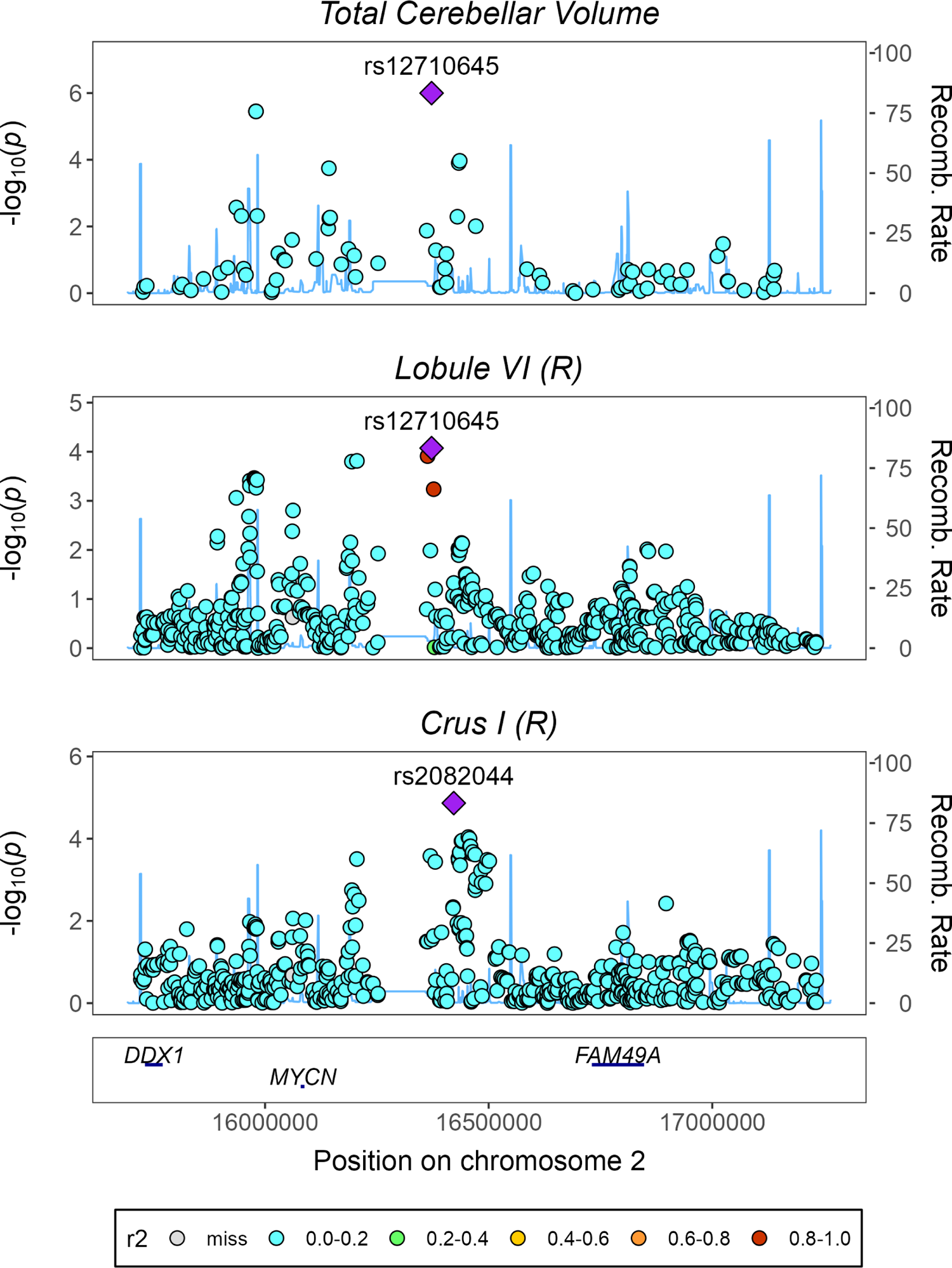
Regional association plot for traits with a significant local genomic correlation in locus 2:15721704-17241115, (LAVA). HapMap3 SNPs are shown.

**Figure S10.**
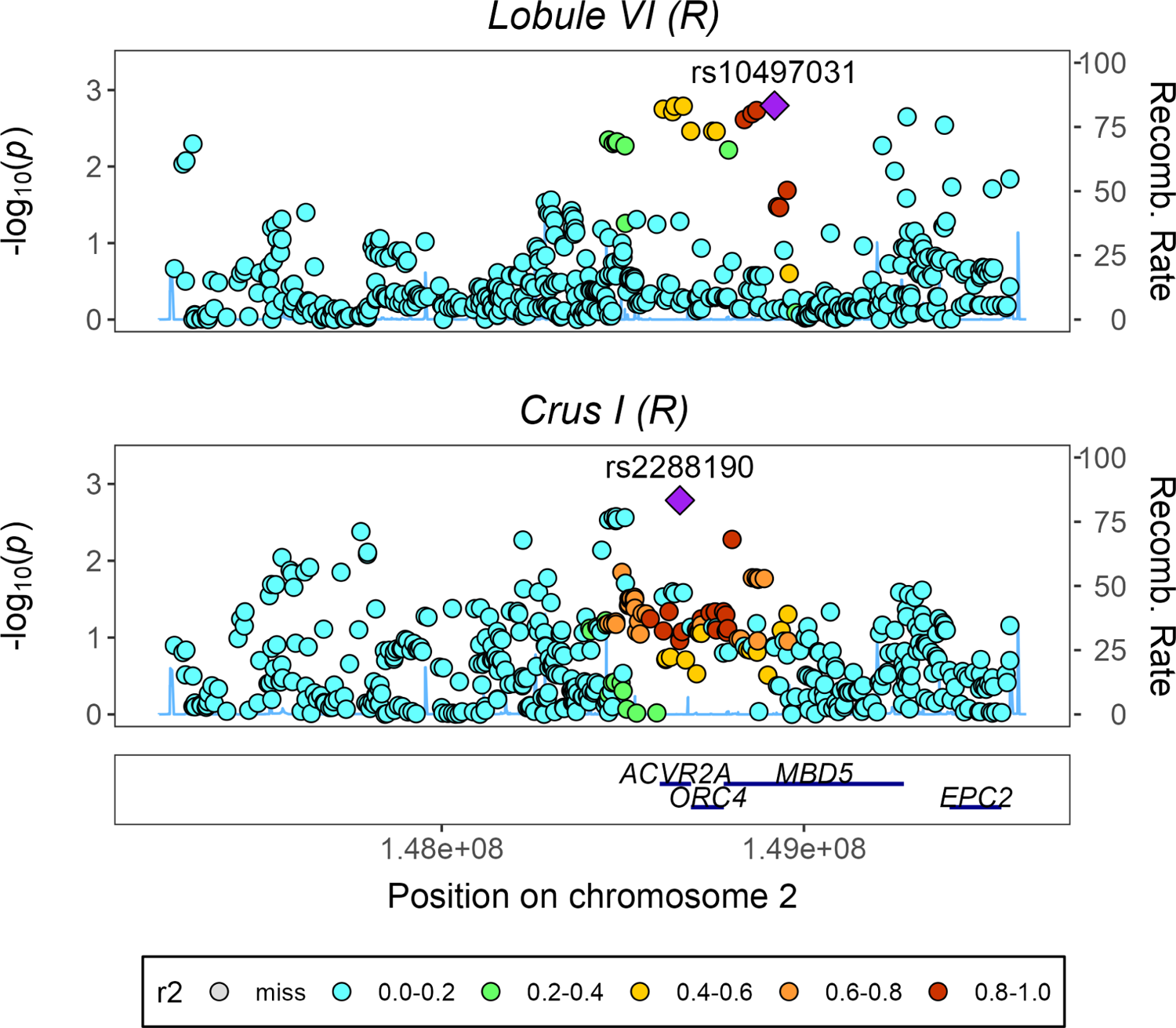
Regional association plot for traits with a significant local genomic correlation in locus 2:147252197-149568875, (LAVA). HapMap3 SNPs are shown.

**Figure S11.**
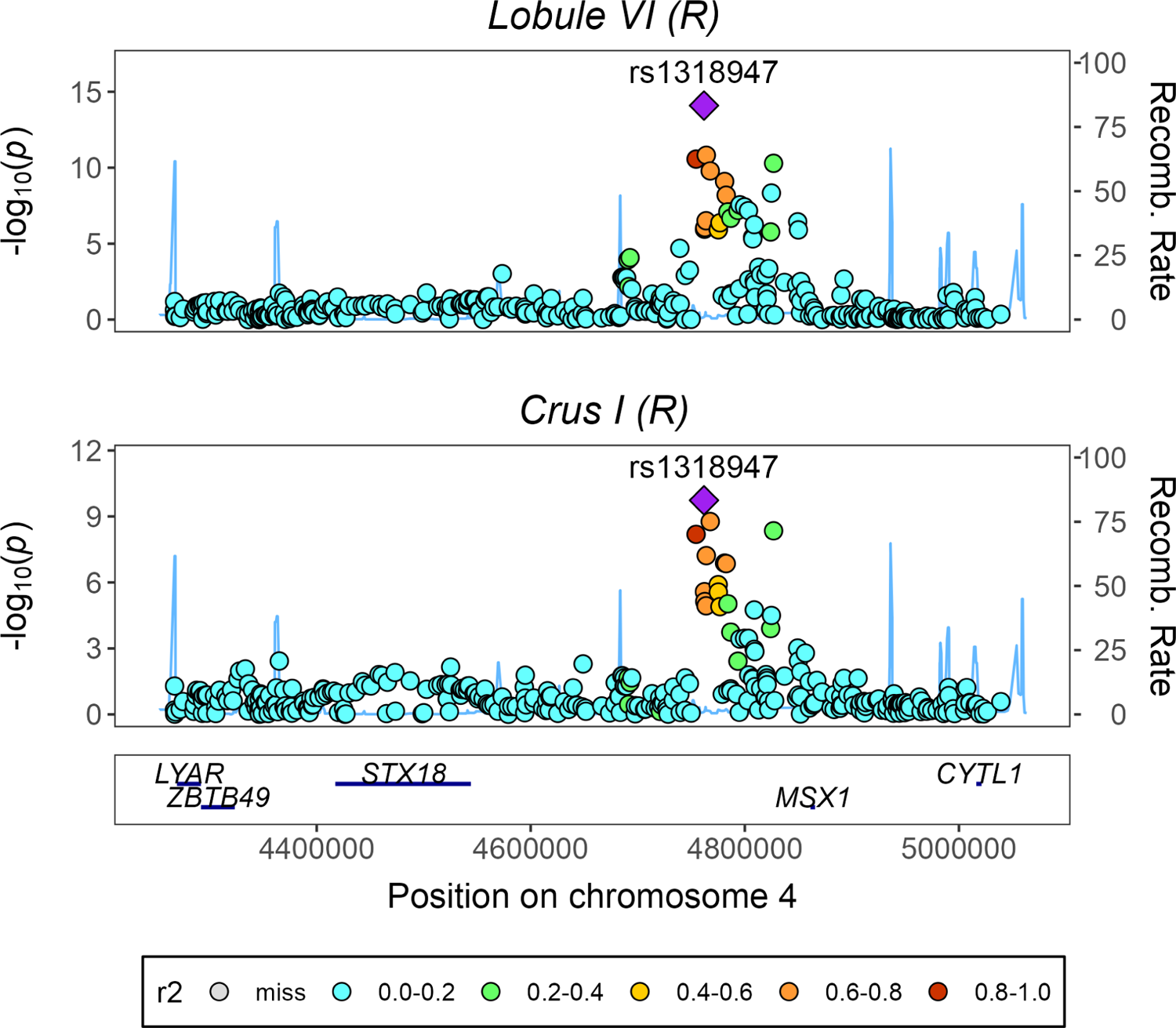
Regional association plot for traits with a significant local genomic correlation in locus 4:4266179-5051834, (LAVA). HapMap3 SNPs are shown.

**Figure S12.**
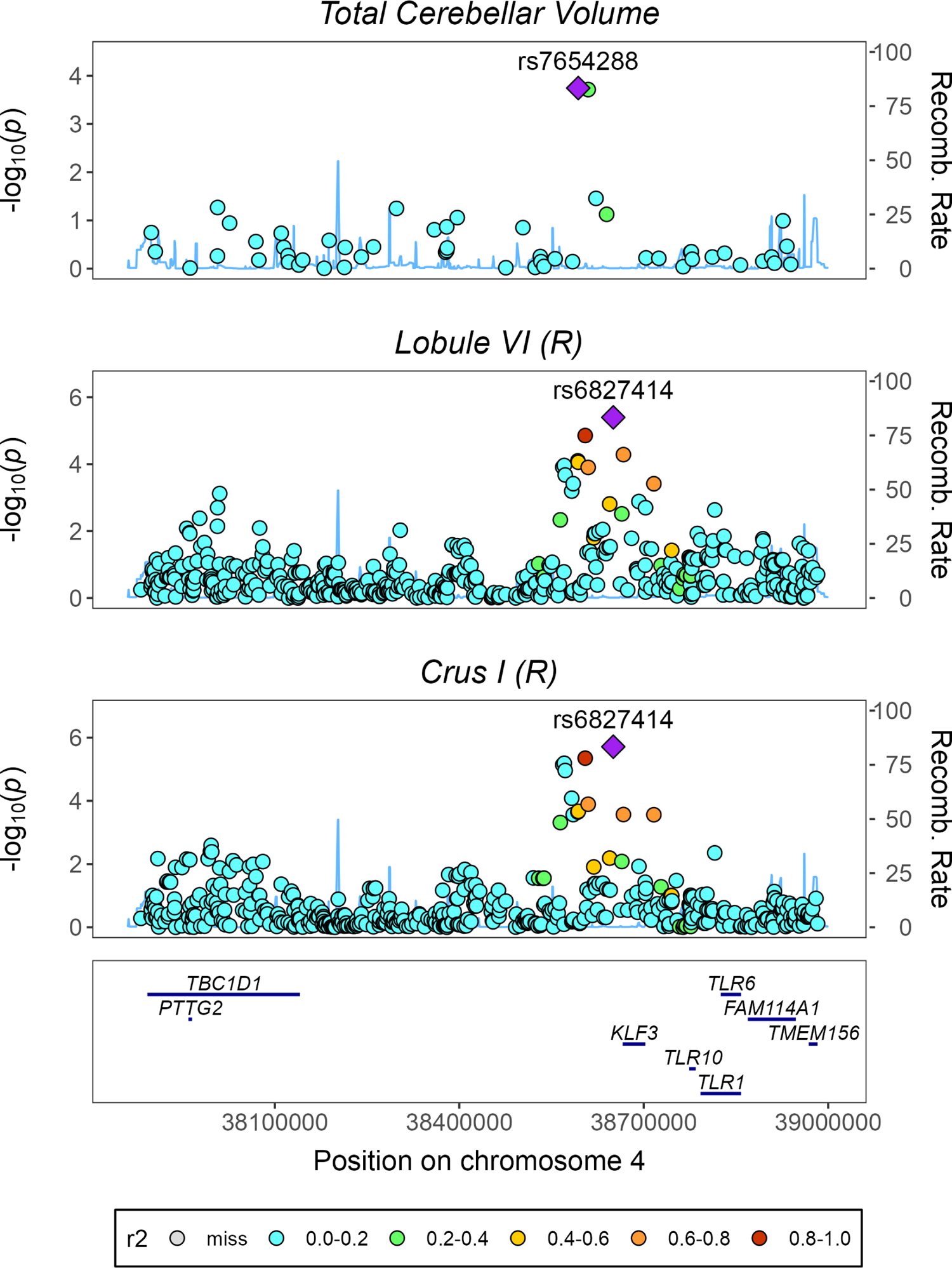
Regional association plot for traits with a significant local genomic correlation in locus 4:37880861-38984838, (LAVA). HapMap3 SNPs are shown.

**Figure S13.**
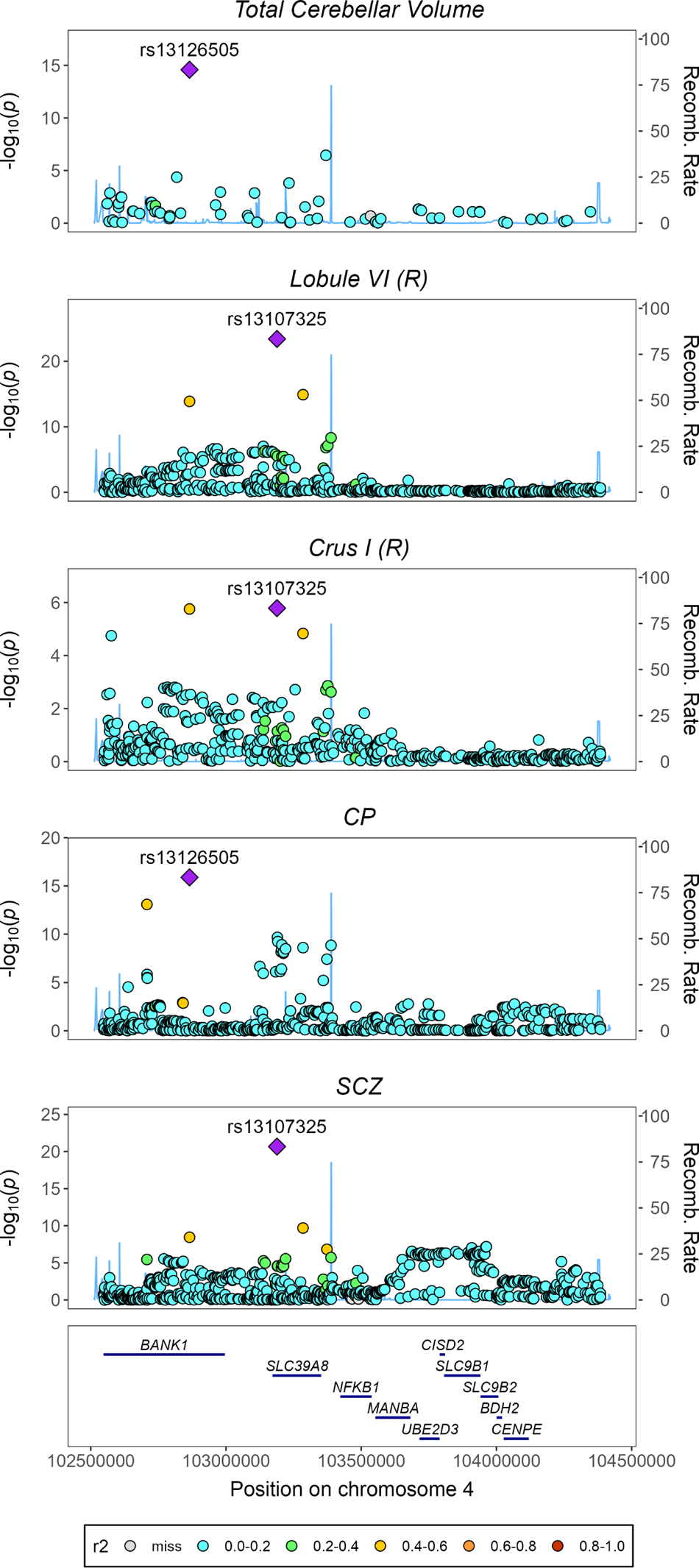
Regional association plot for traits with a significant local genomic correlation in locus 4:102544804-104384534, (LAVA). HapMap3 SNPs are shown.

**Figure S14.**
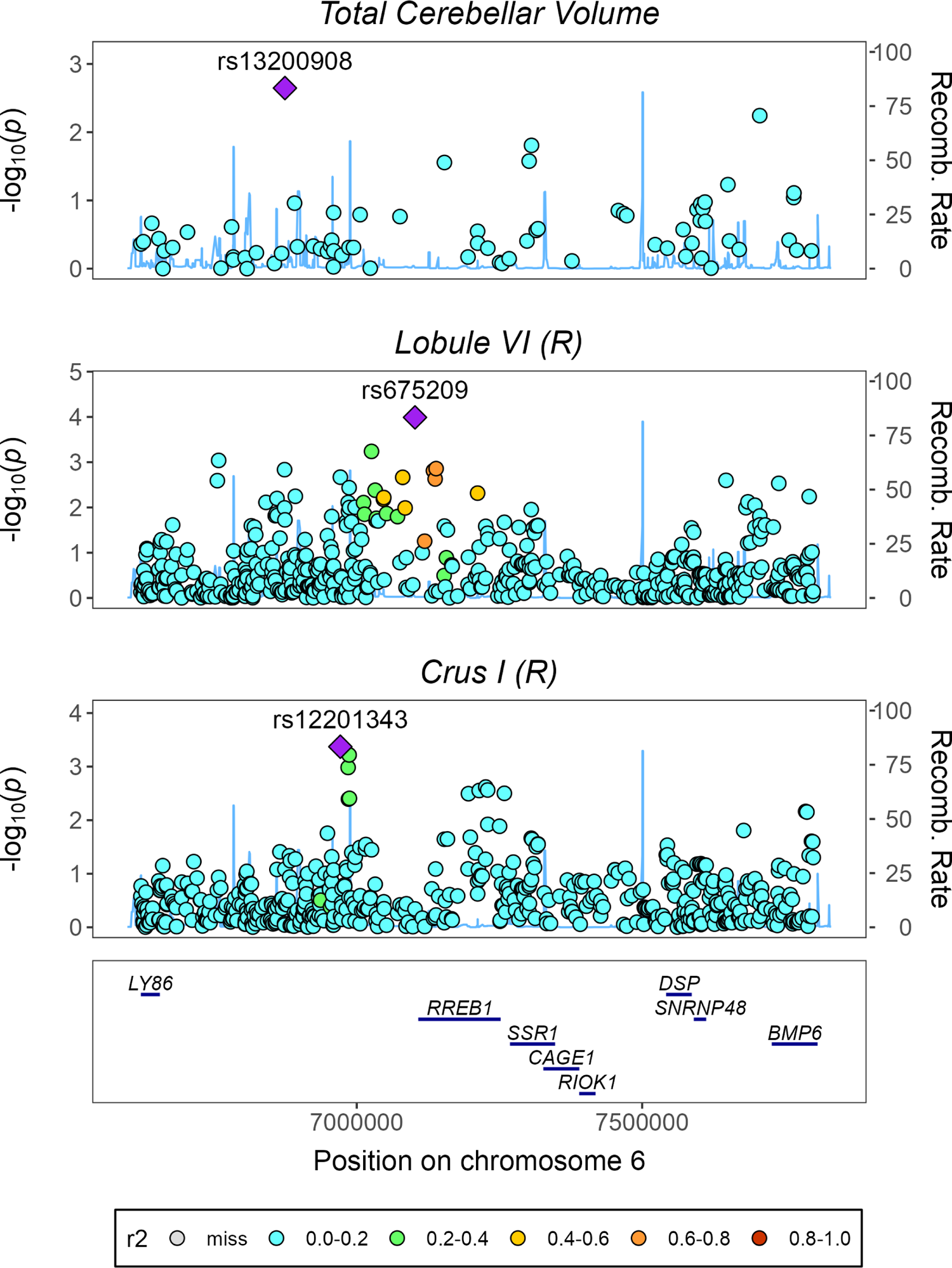
Regional association plot for traits with a significant local genomic correlation in locus 6:6621938-7808323, (LAVA). HapMap3 SNPs are shown.

**Figure S15.**
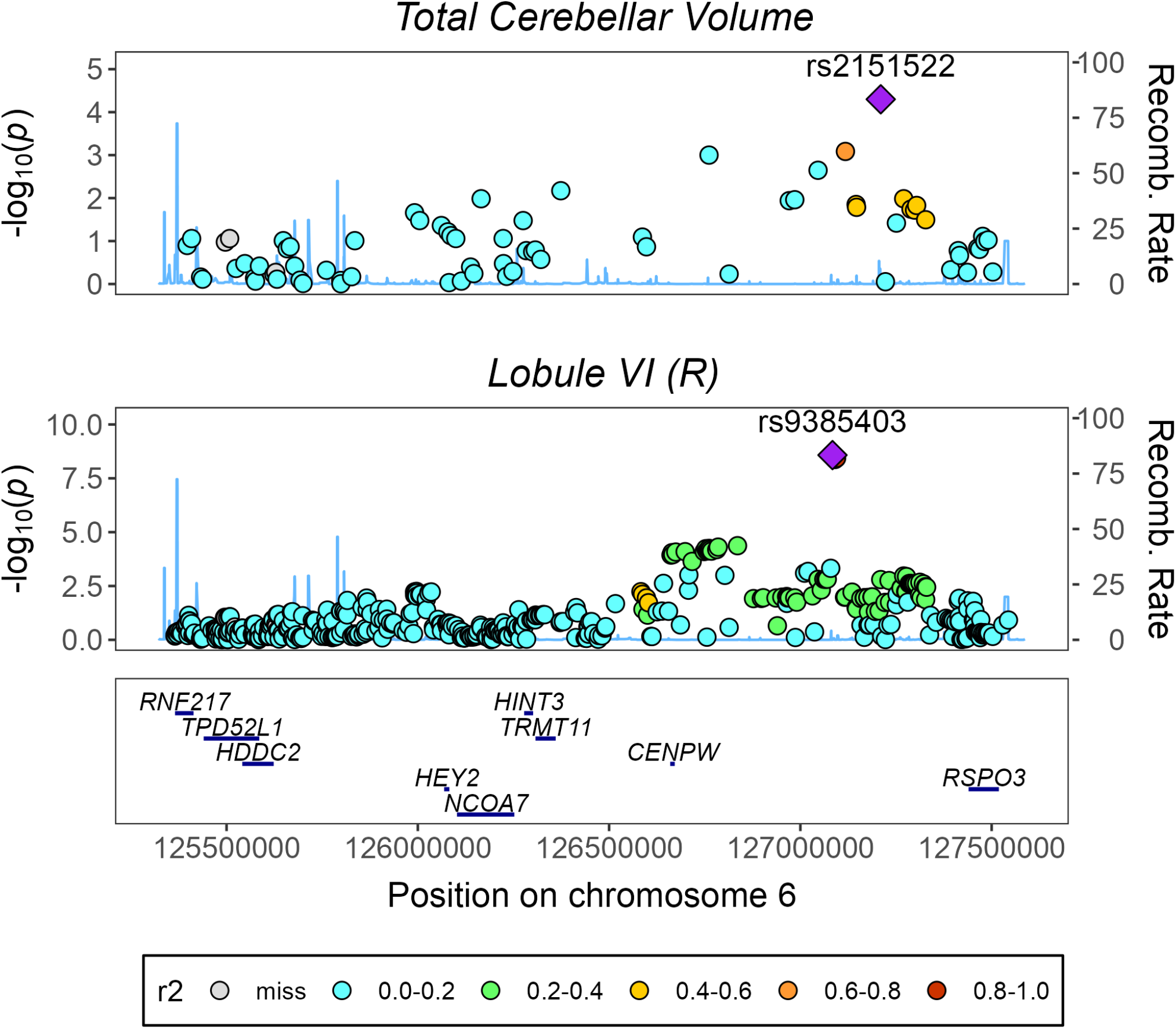
Regional association plot for traits with a significant local genomic correlation in locus 6:125365055-127545459, (LAVA). HapMap3 SNPs are shown.

**Figure S16.**
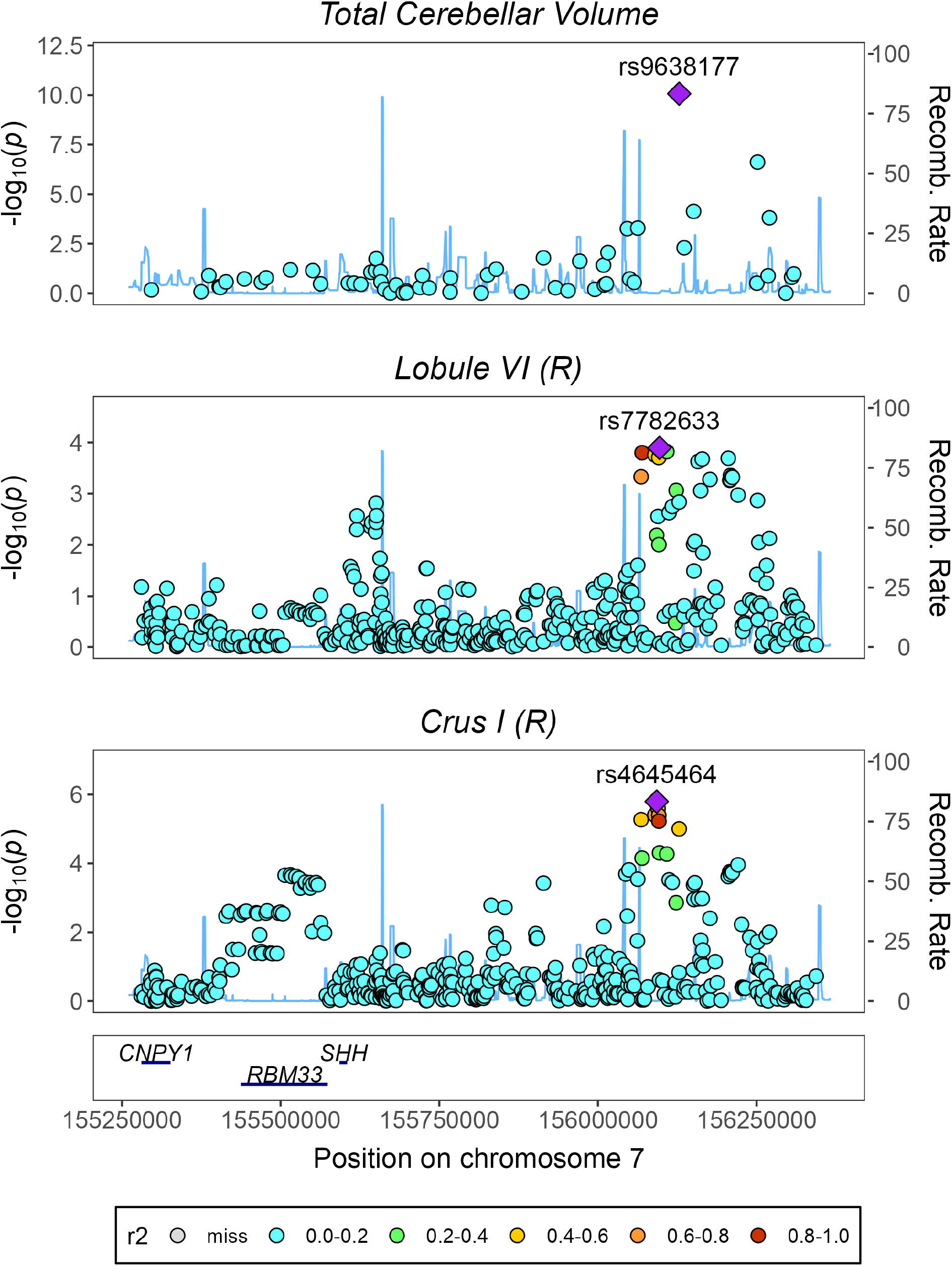
Regional association plot for traits with a significant local genomic correlation in locus 7:155280611-156344386, (LAVA). HapMap3 SNPs are shown.

**Figure S17.**
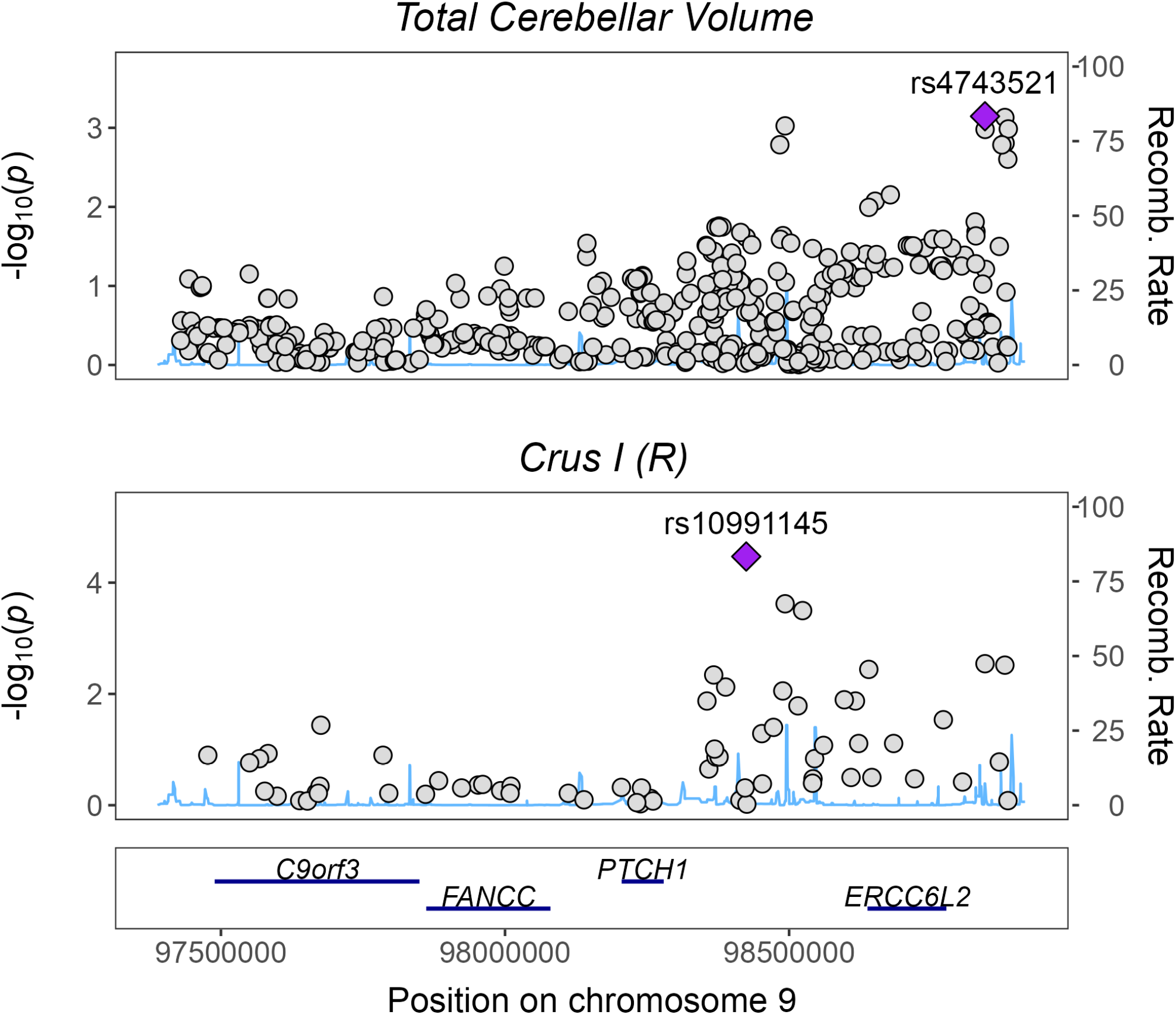
Regional association plot for traits with a significant local genomic correlation in locus 9:97418634-98885862, (LAVA). HapMap3 SNPs are shown.

**Figure S18.**
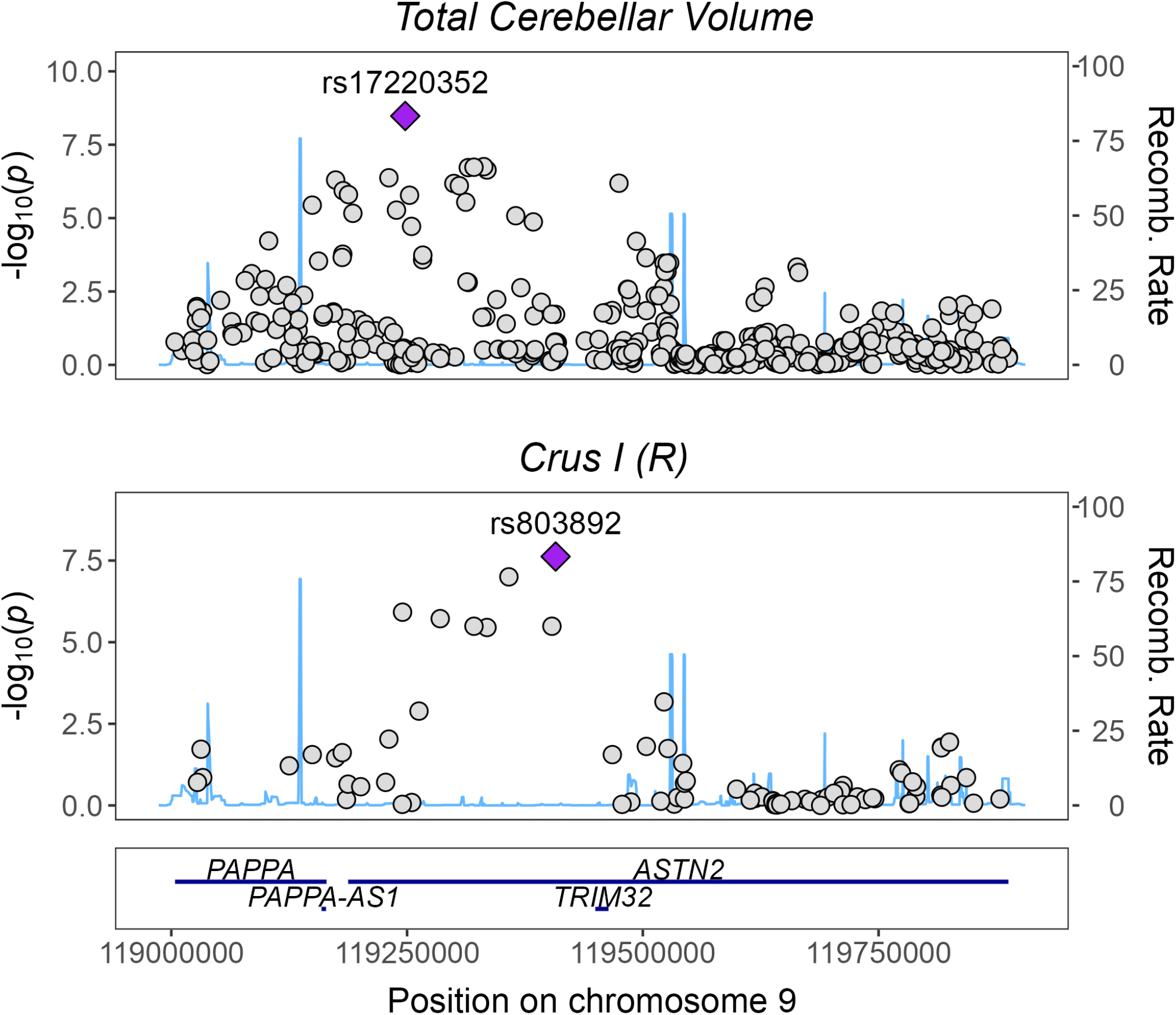
Regional association plot for traits with a significant local genomic correlation in locus 9:119001228-119890464, (LAVA). HapMap3 SNPs are shown.

**Figure S19.**
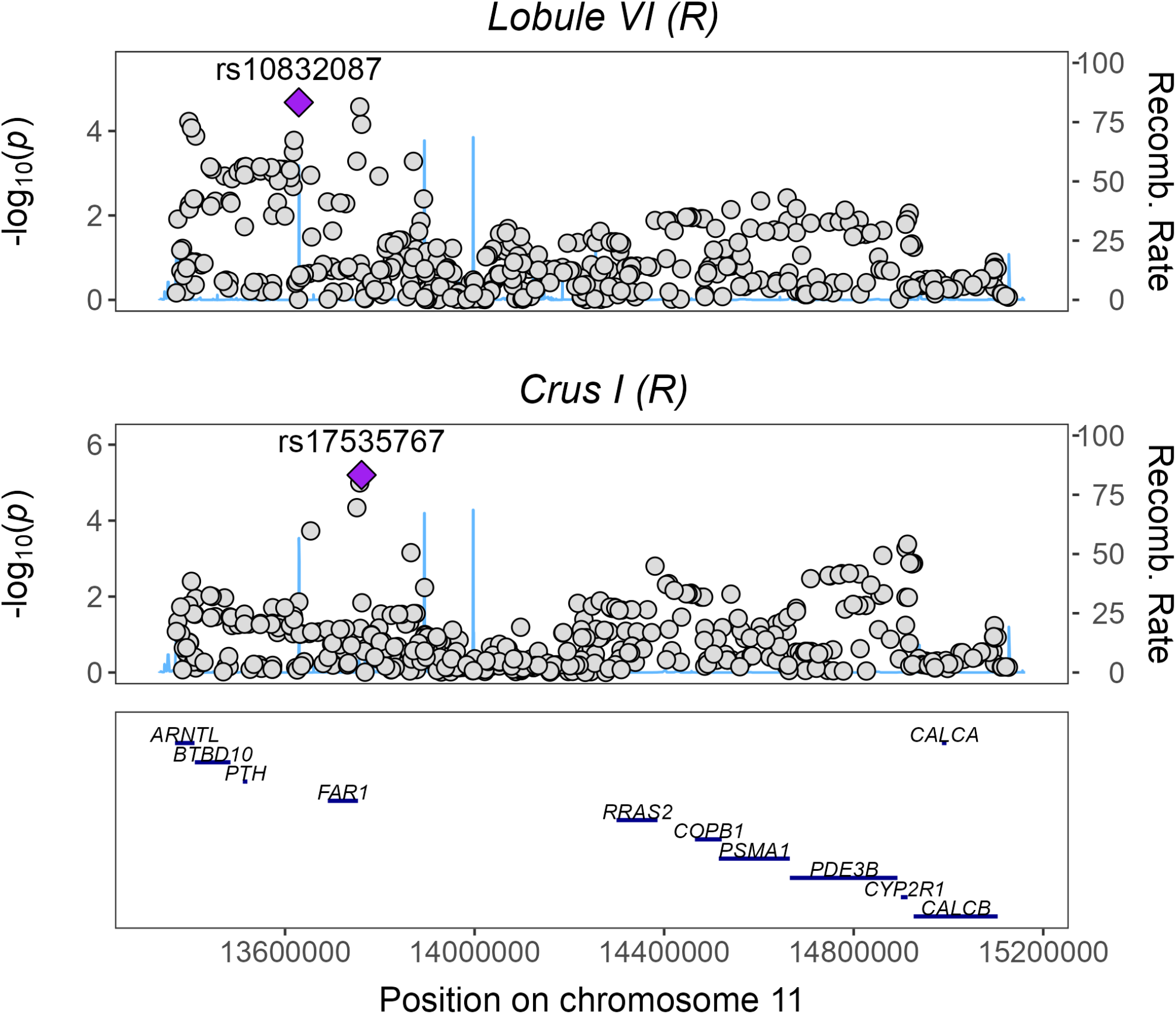
Regional association plot for traits with a significant local genomic correlation in locus 11:13364728-15126767, (LAVA). HapMap3 SNPs are shown.

**Figure S20.**
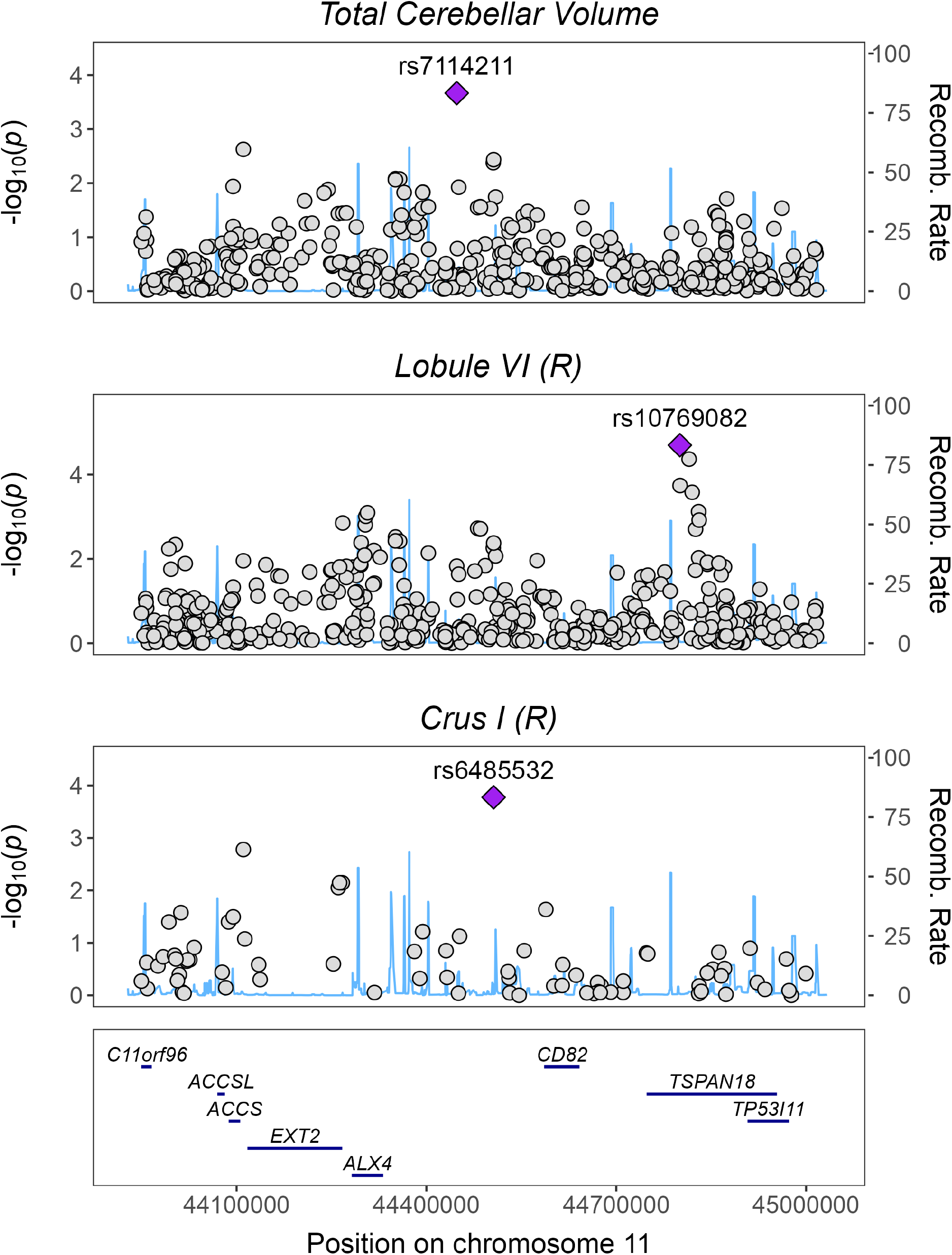
Regional association plot for traits with a significant local genomic correlation in locus 11:43949378-45019559, (LAVA). HapMap3 SNPs are shown.

**Figure S21.**
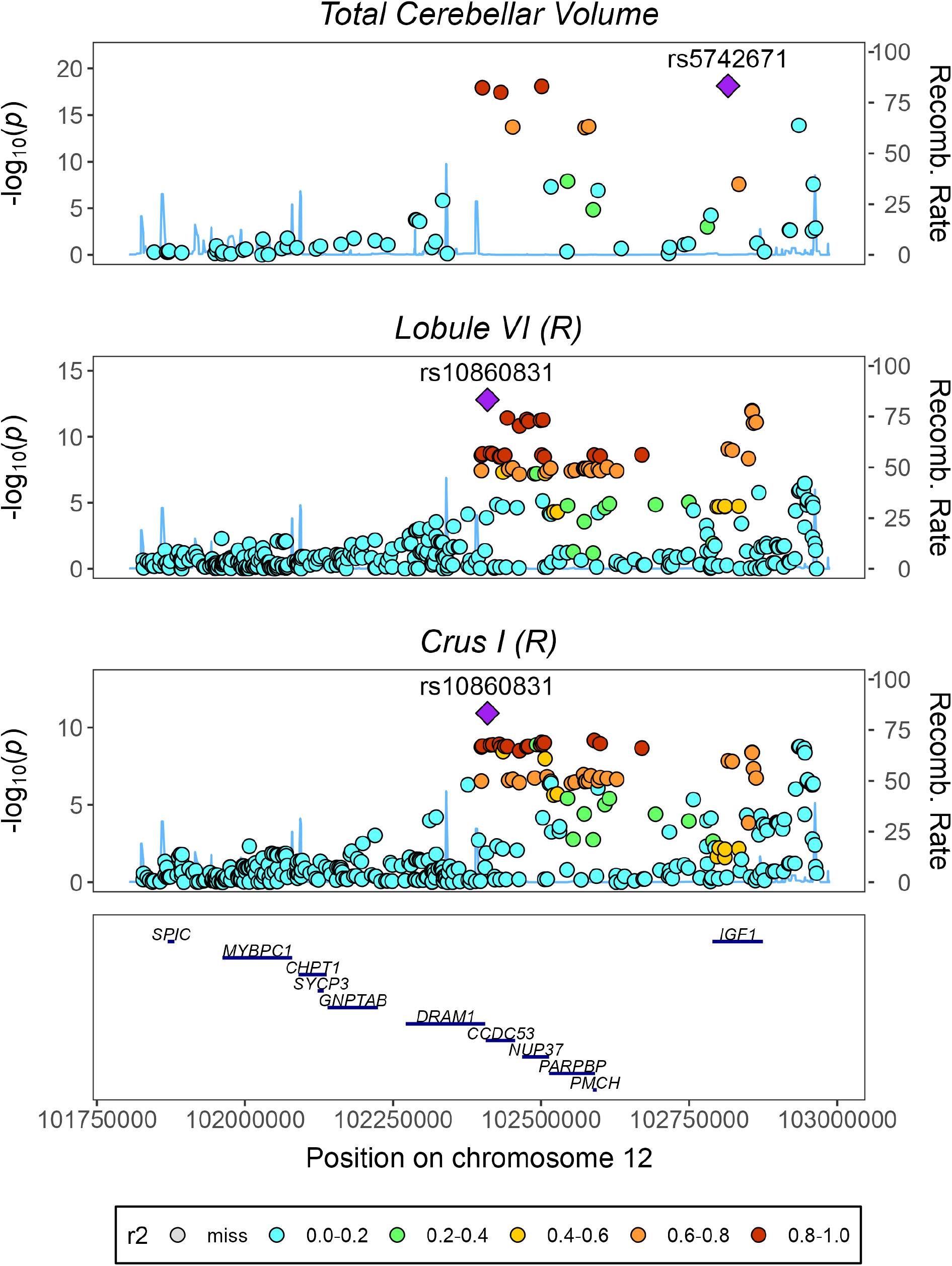
Regional association plot for traits with a significant local genomic correlation in locus 12:101825062-102966633, (LAVA). HapMap3 SNPs are shown.

**Figure S22.**
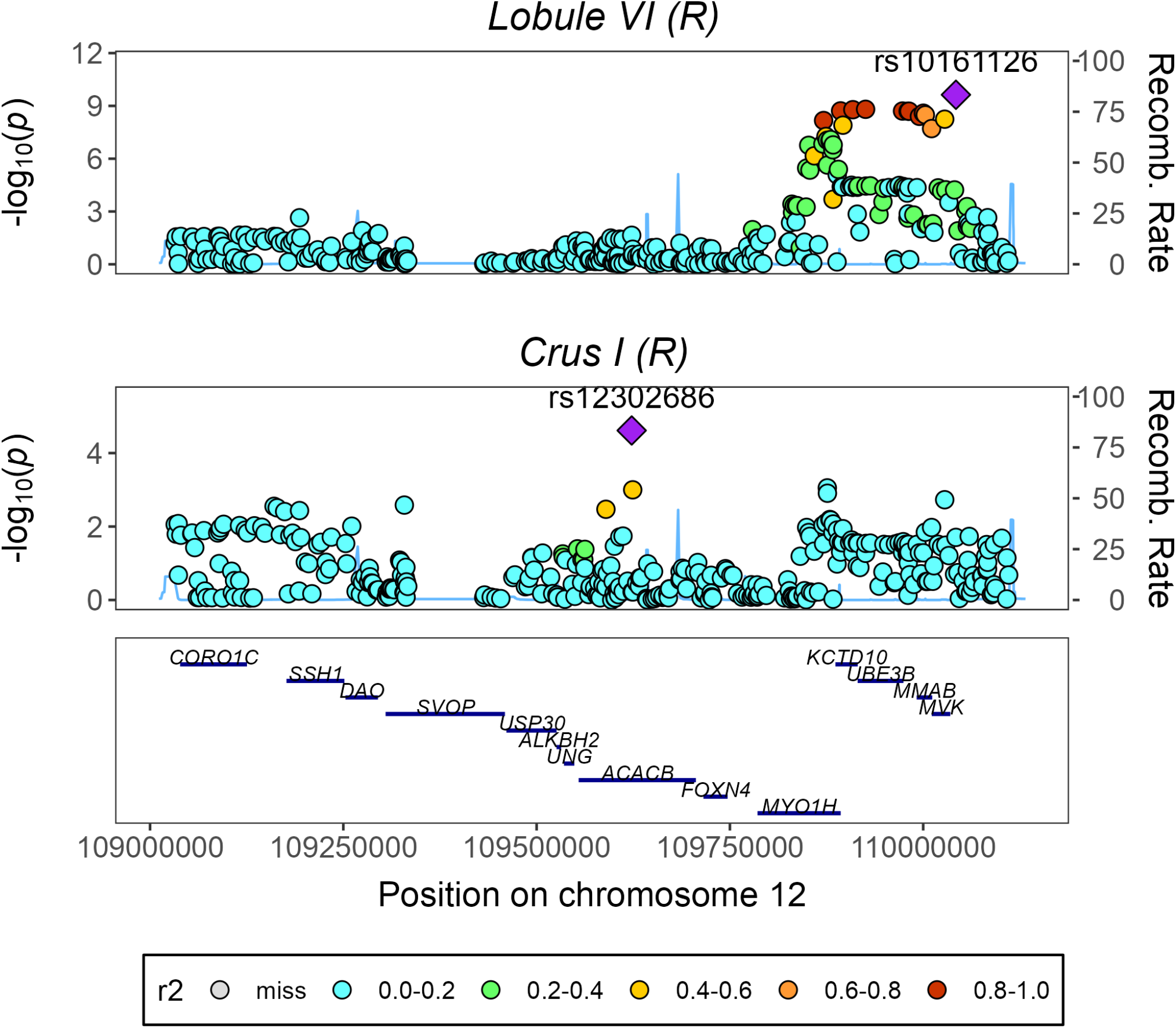
Regional association plot for traits with a significant local genomic correlation in locus 12:109031820-110111514, (LAVA). HapMap3 SNPs are shown.

**Figure S23.**
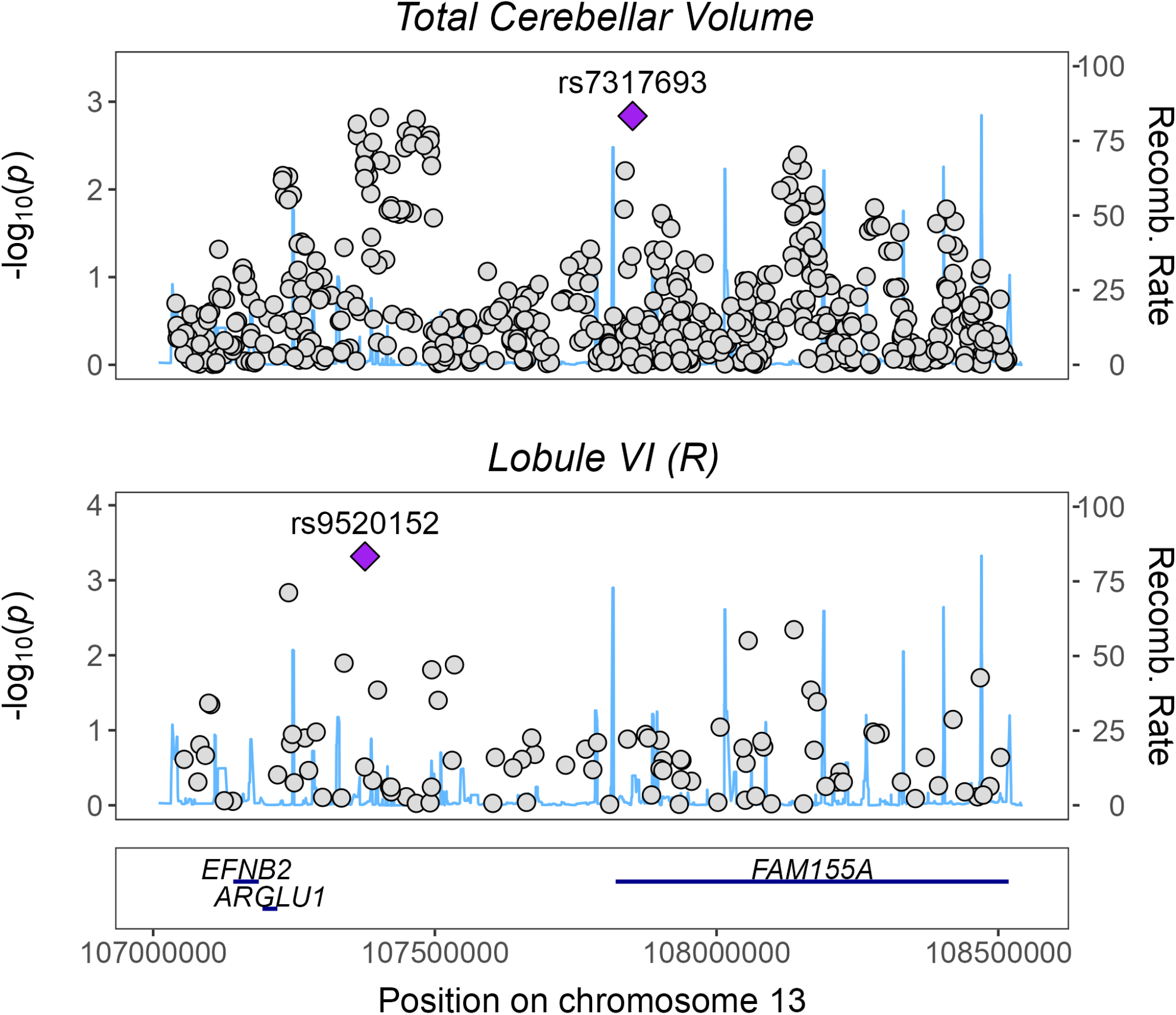
Regional association plot for traits with a significant local genomic correlation in locus 13:107037865-108521978, (LAVA). HapMap3 SNPs are shown.

**Figure S24.**
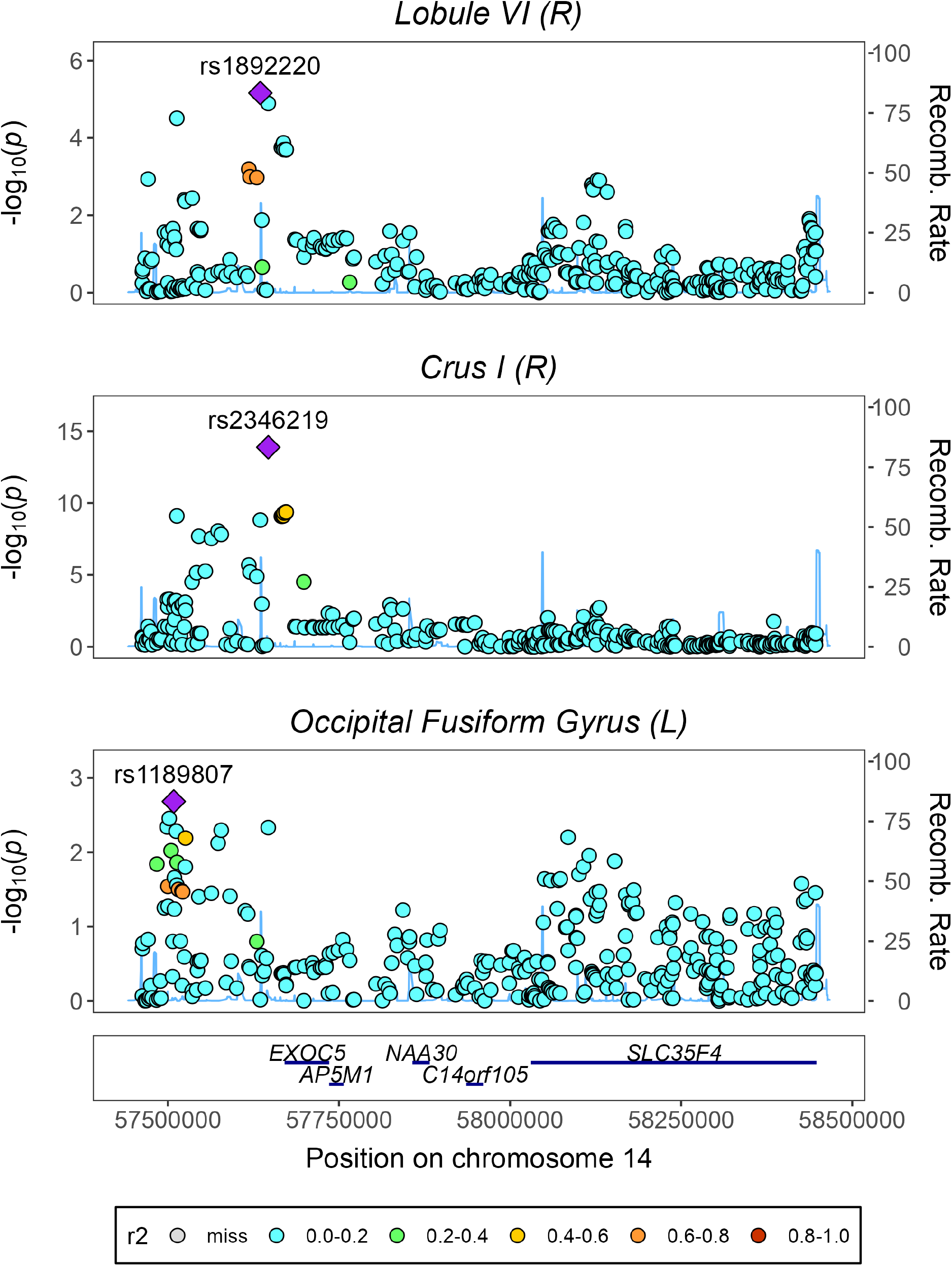
Regional association plot for traits with a significant local genomic correlation in locus 14:57460782-58447798, (LAVA). HapMap3 SNPs are shown.

**Figure S25.**
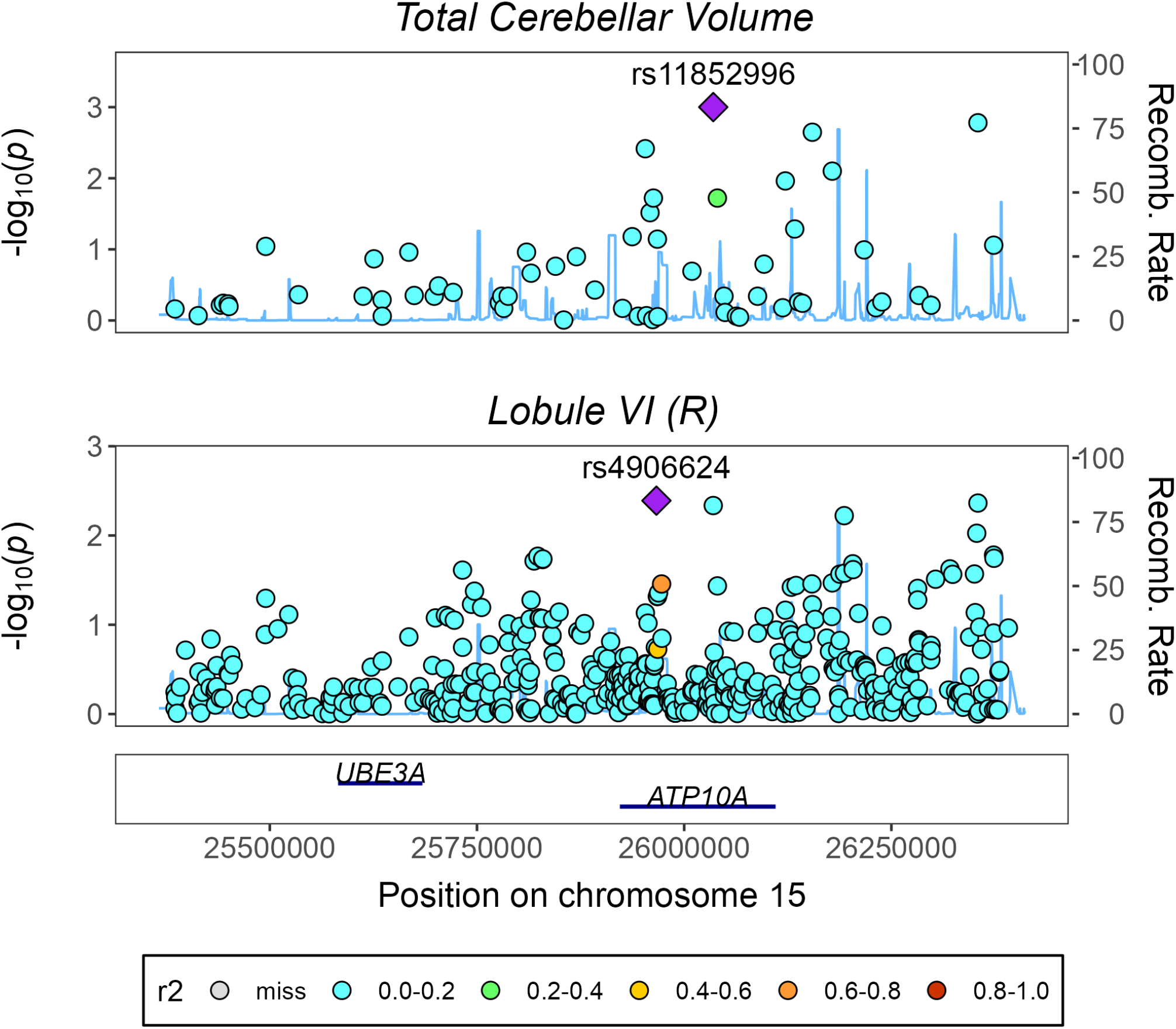
Regional association plot for traits with a significant local genomic correlation in locus 15:25384328-26392947, (LAVA). HapMap3 SNPs are shown.

**Figure S26.**
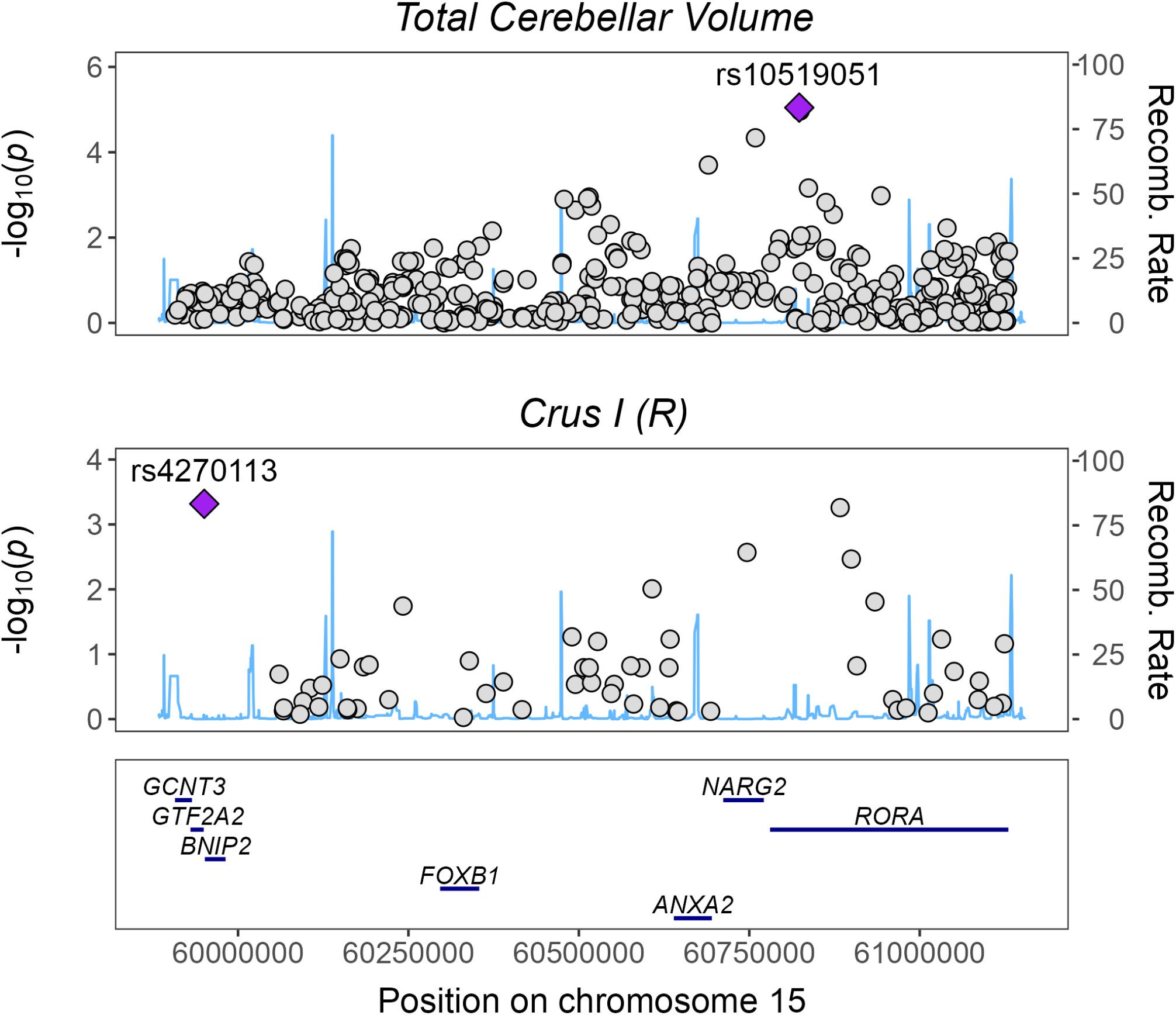
Regional association plot for traits with a significant local genomic correlation in locus 15:59901117-61130595, (LAVA). HapMap3 SNPs are shown.

**Figure S27.**
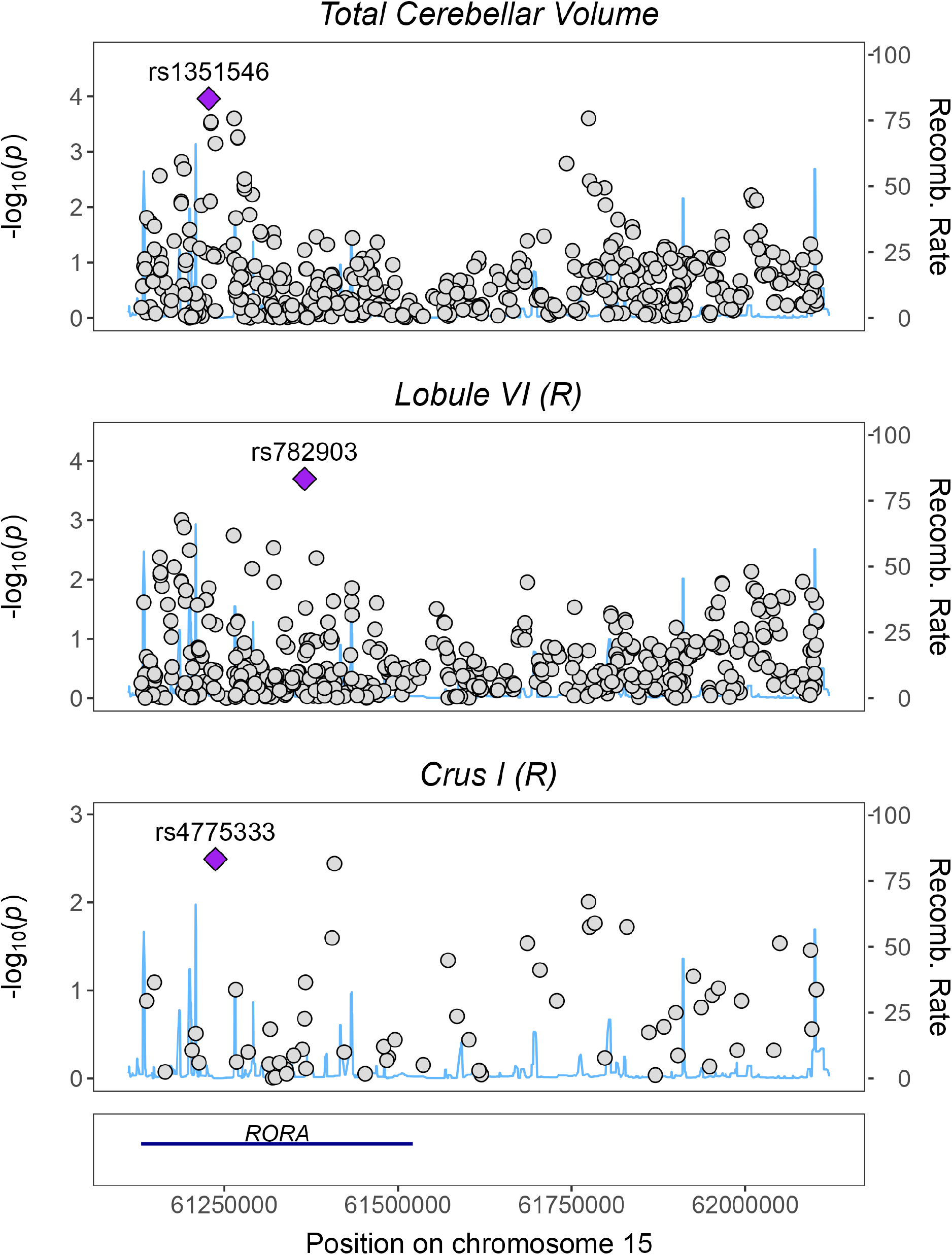
Regional association plot for traits with a significant local genomic correlation in locus 15:61130596-62106138, (LAVA). HapMap3 SNPs are shown.

**Figure S28.**
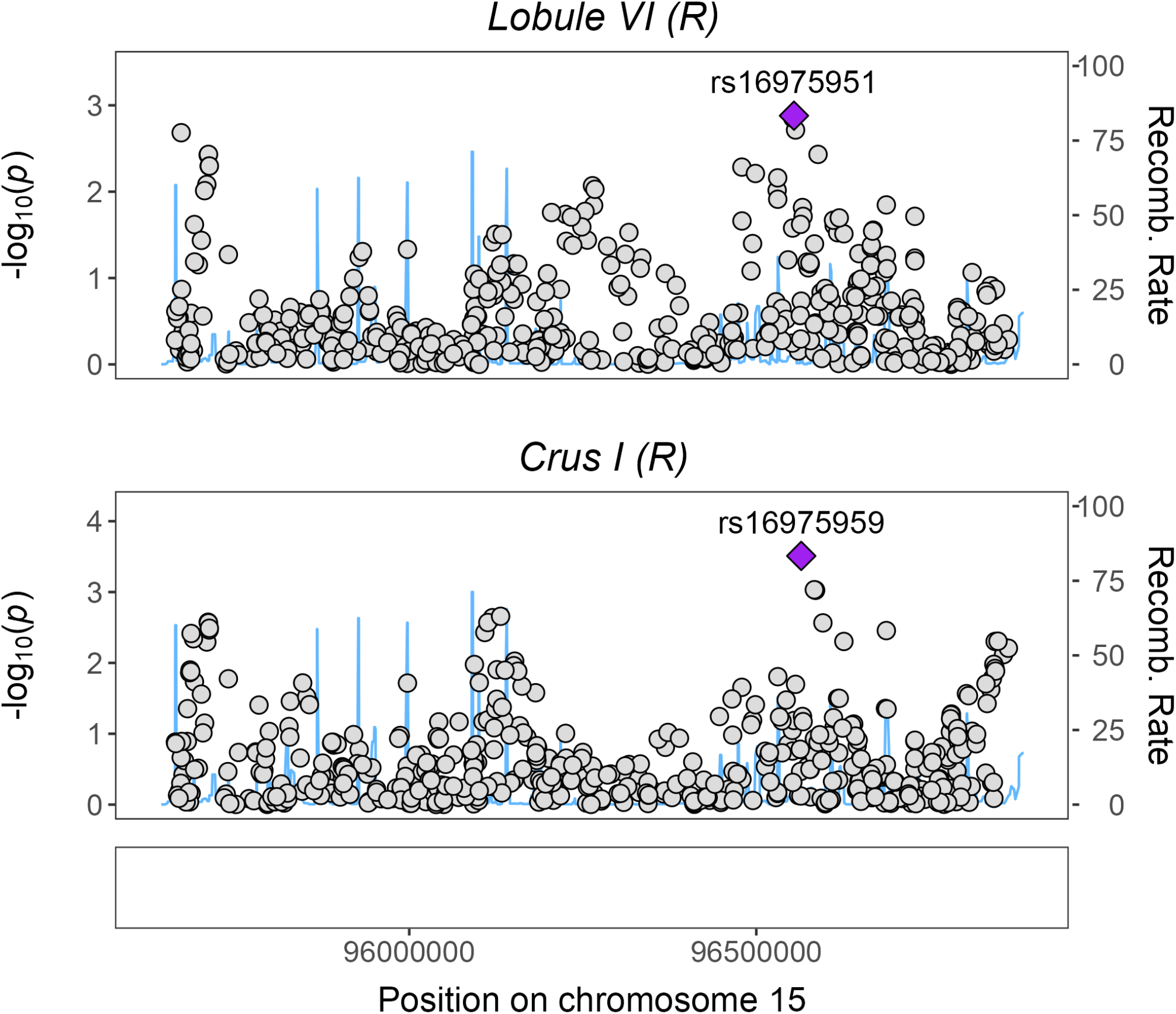
Regional association plot for traits with a significant local genomic correlation in locus 15:95662426-96864278, (LAVA). HapMap3 SNPs are shown.

**Figure S29.**
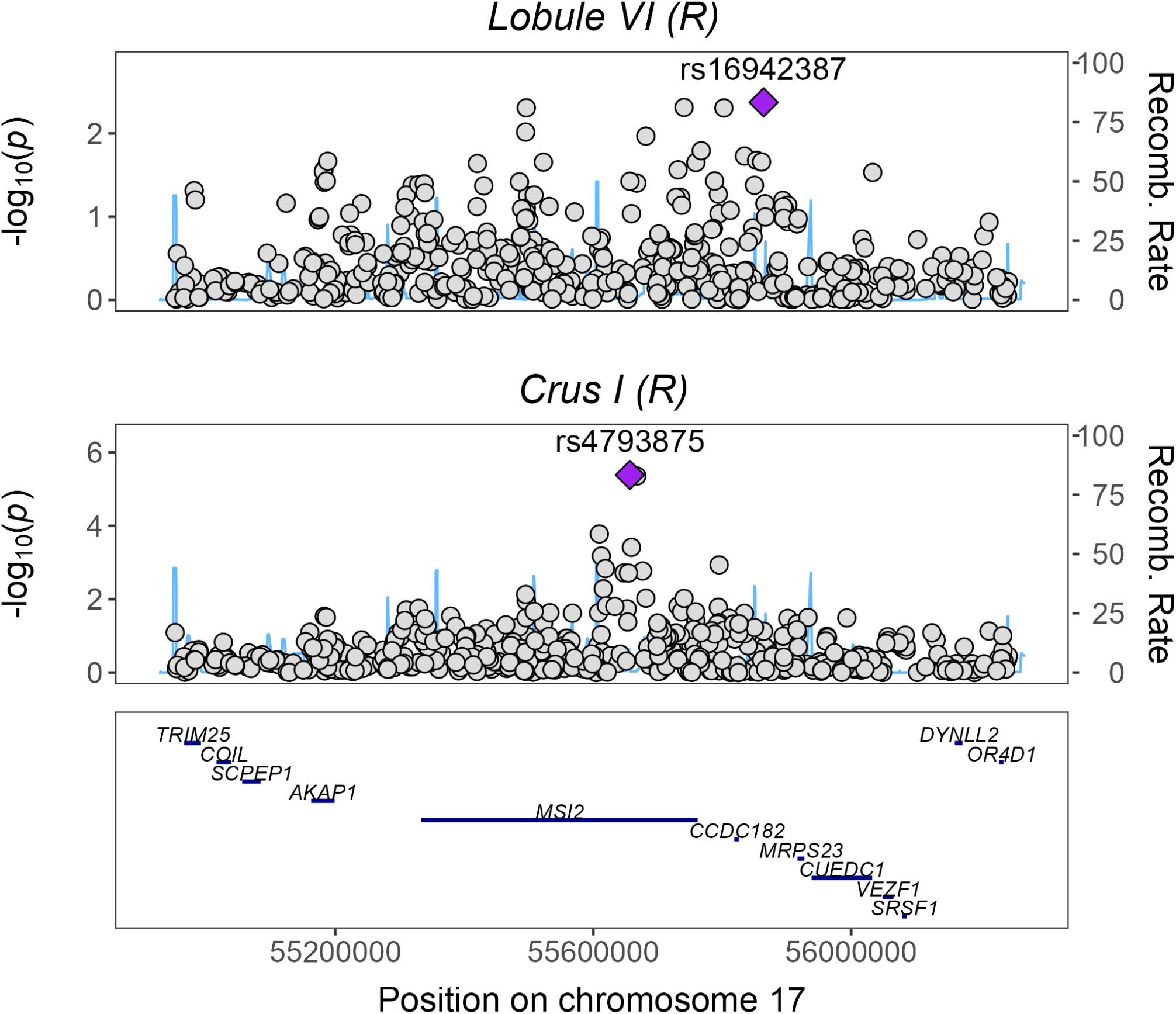
Regional association plot for traits with a significant local genomic correlation in locus 17:54950108-56245227, (LAVA). HapMap3 SNPs are shown.

**Figure S30.**
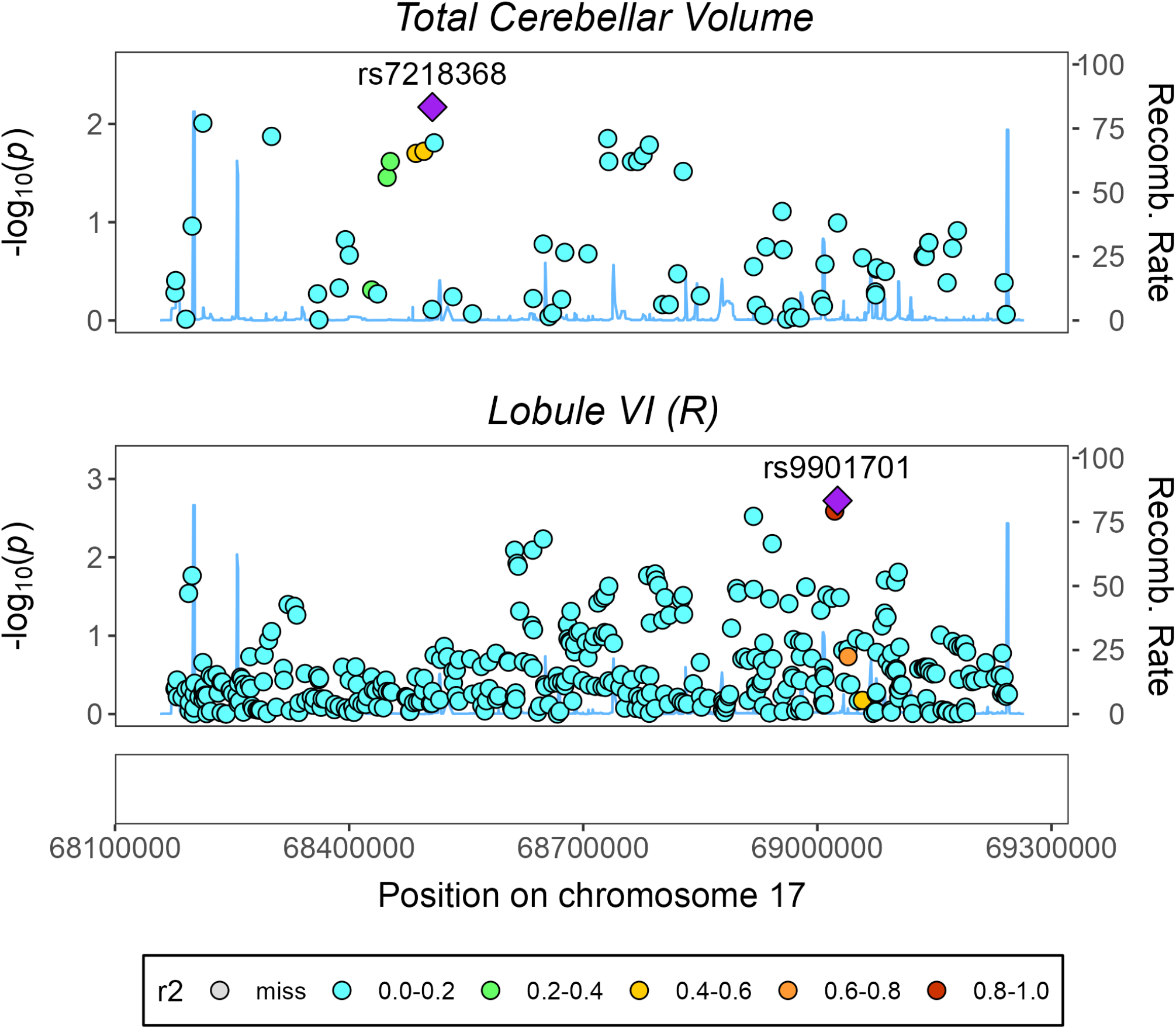
Regional association plot for traits with a significant local genomic correlation in locus 17:68176220-69245590, (LAVA). HapMap3 SNPs are shown.

**Figure S31.**
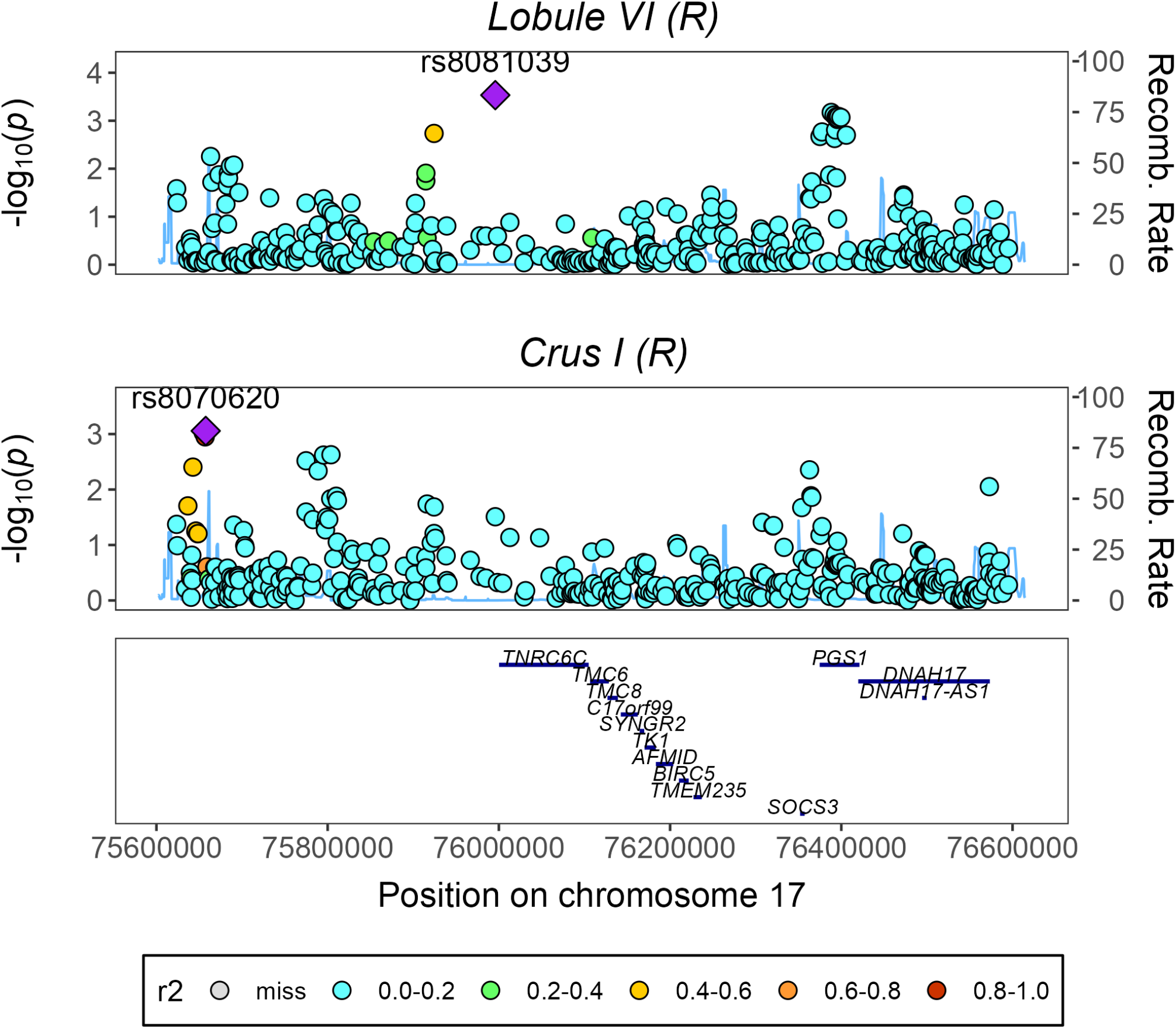
Regional association plot for traits with a significant local genomic correlation in locus 17:75619257-76596287, (LAVA). HapMap3 SNPs are shown.

**Figure S32.**
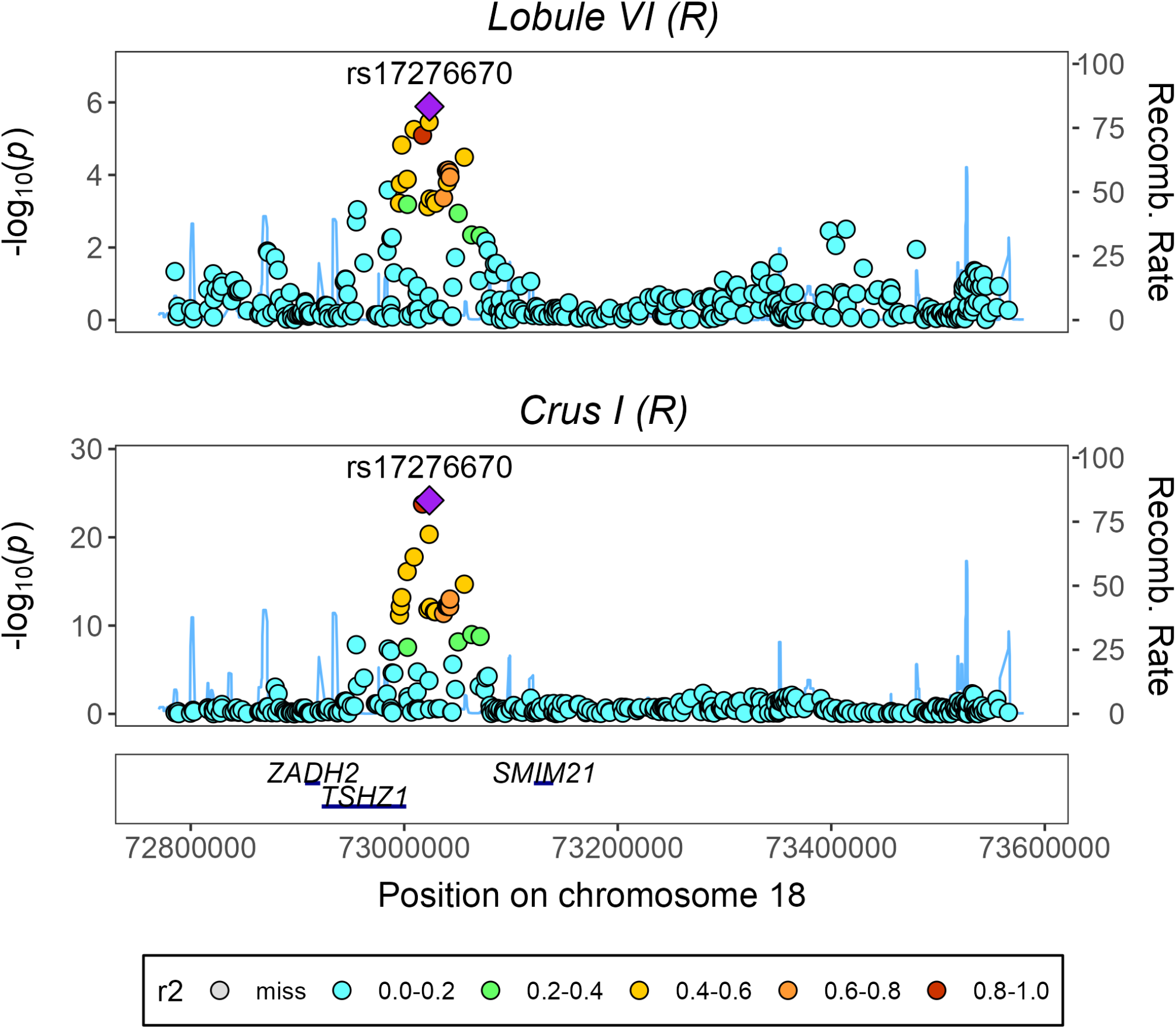
Regional association plot for traits with a significant local genomic correlation in locus 18:72785207-73568260, (LAVA). HapMap3 SNPs are shown.

**Figure S33.**
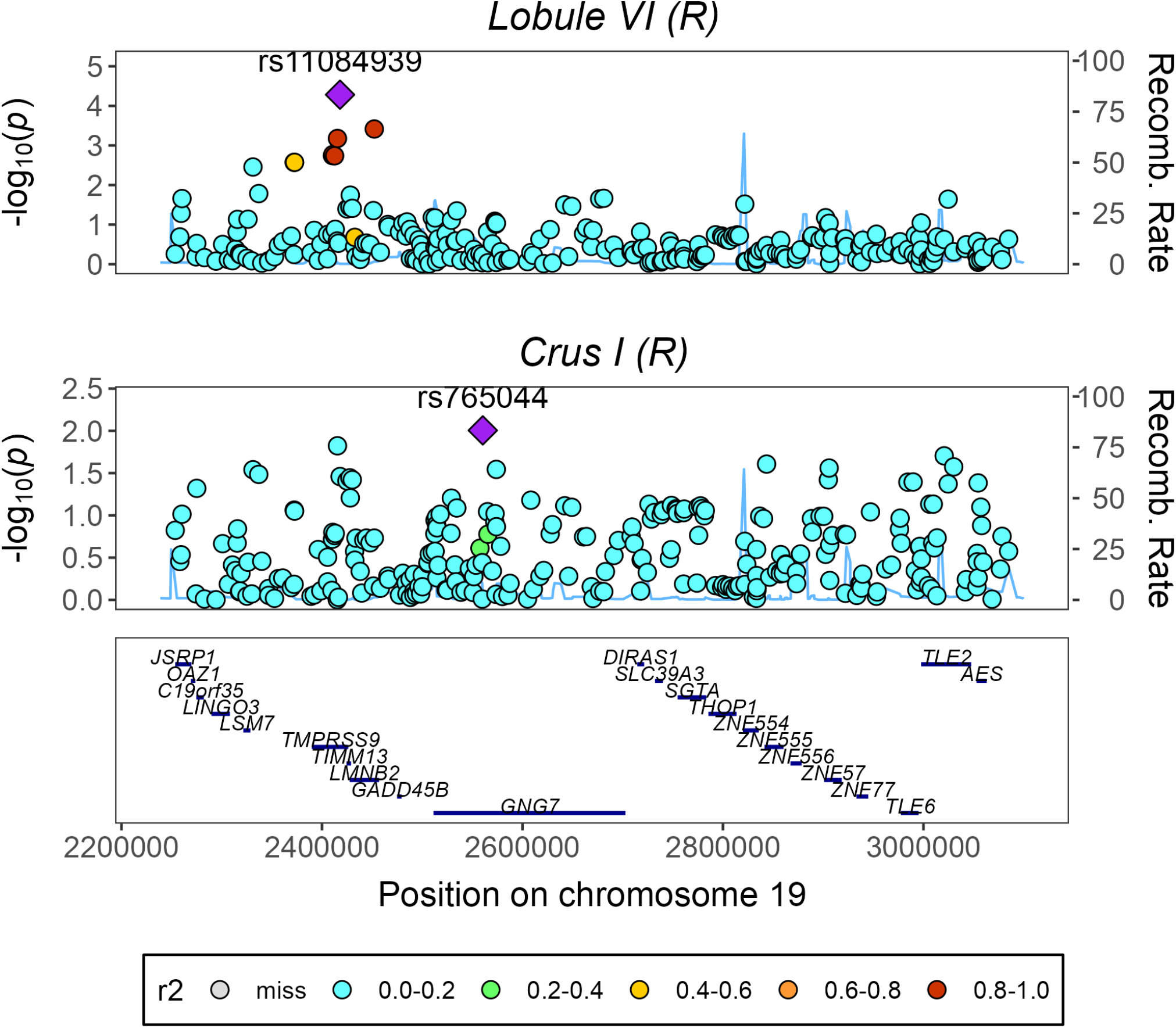
Regional association plot for traits with a significant local genomic correlation in locus 19:2253175-3085446, (LAVA). HapMap3 SNPs are shown.

**Figure S34.**
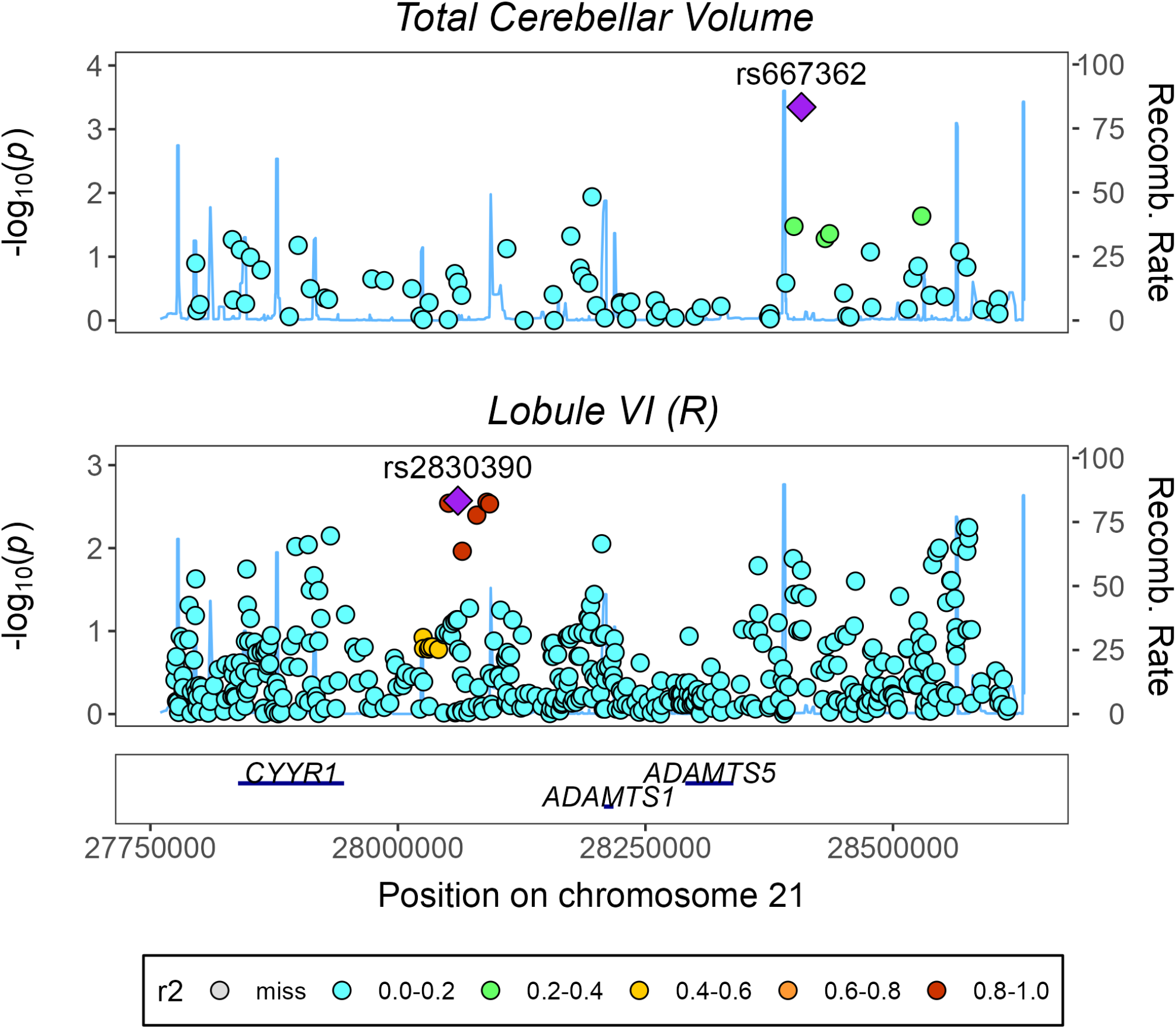
Regional association plot for traits with a significant local genomic correlation in locus 21:27774351-28616539, (LAVA). HapMap3 SNPs are shown.

**Figure S35.**
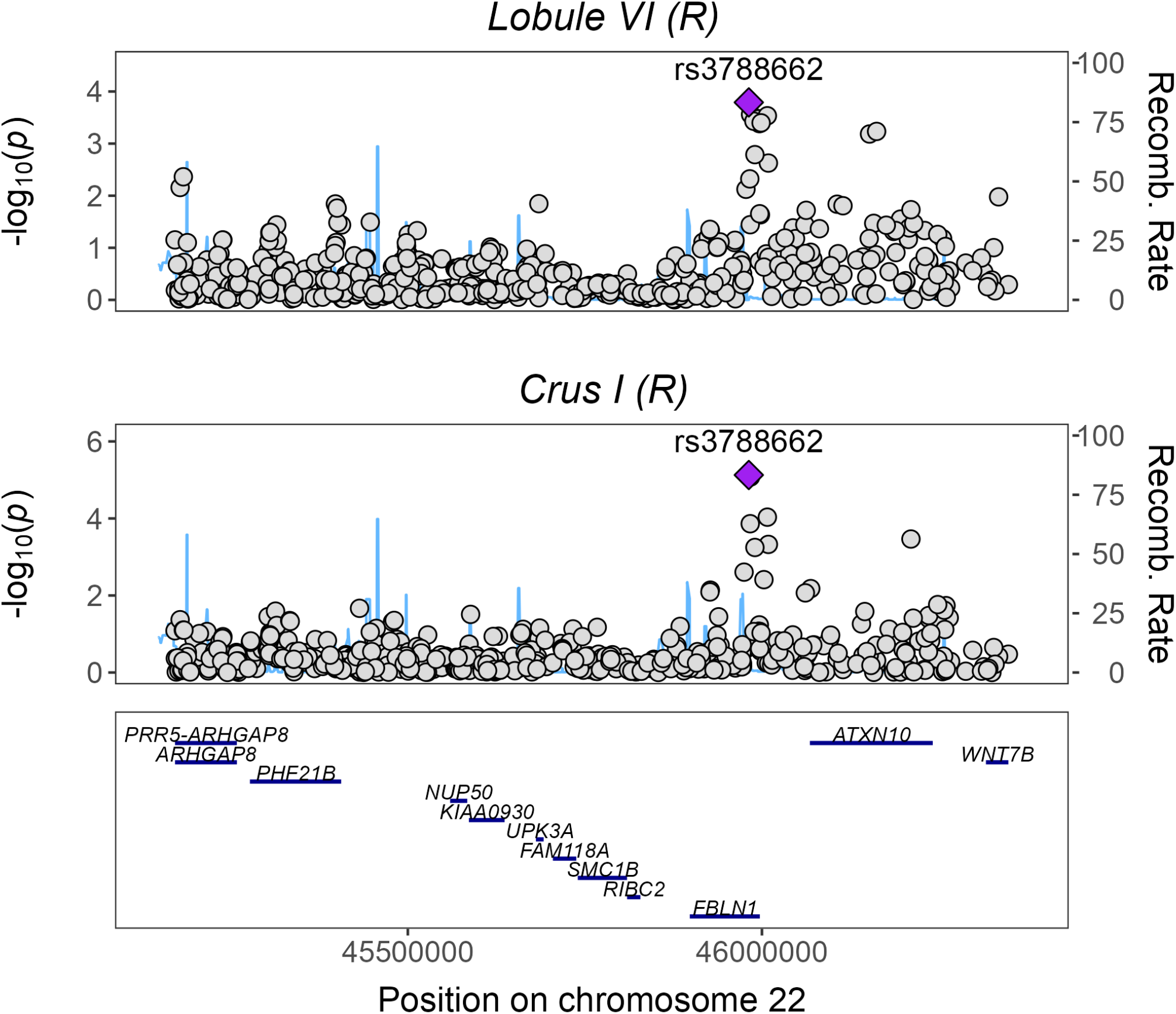
Regional association plot for traits with a significant local genomic correlation in locus 22:45170650-46351048, (LAVA). HapMap3 SNPs are shown.

**Figure S36.**
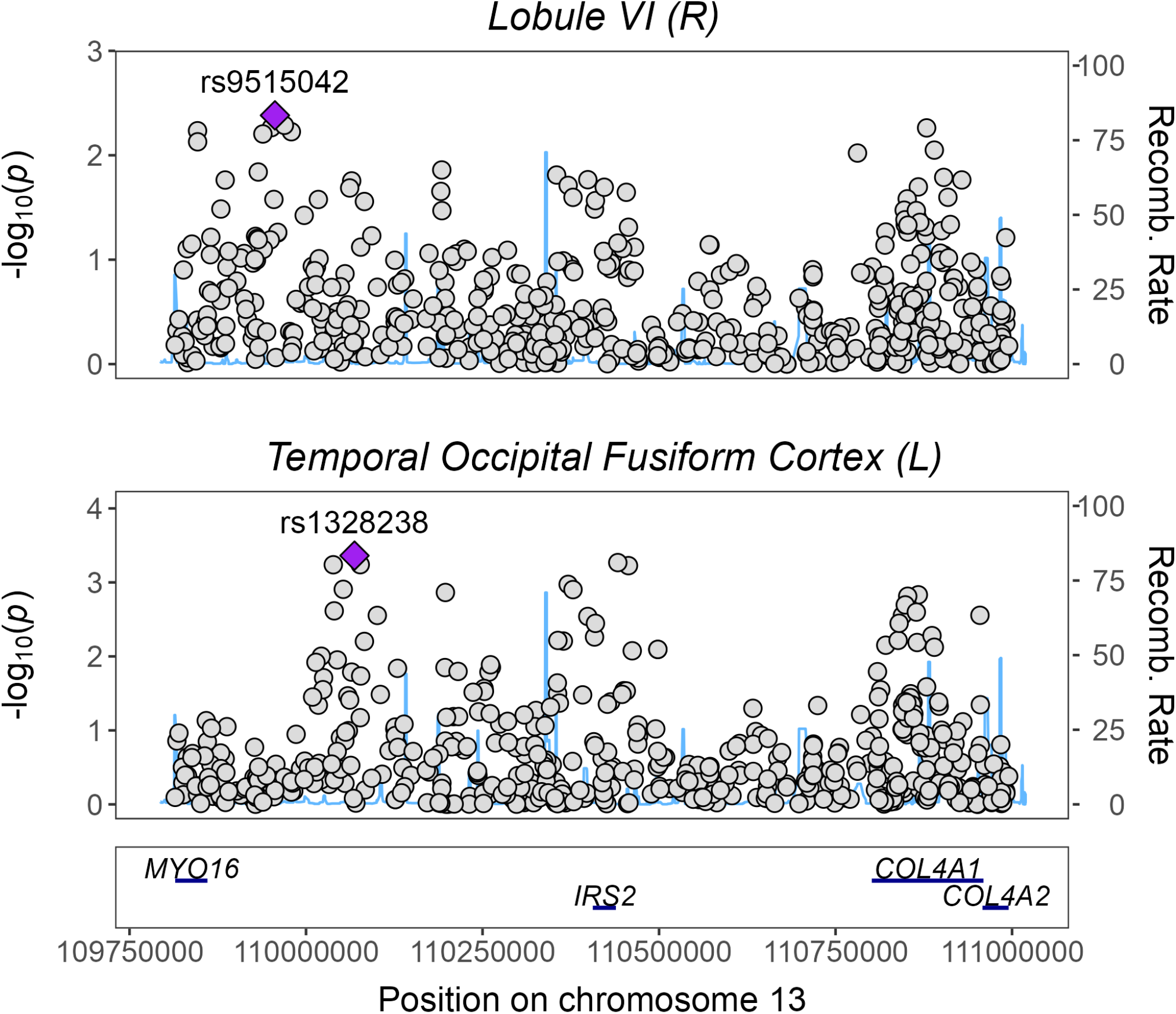
Regional association plot for traits with a significant local genomic correlation in locus 13:109813577-110995432, (LAVA). HapMap3 SNPs are shown.

**Figure S37.**
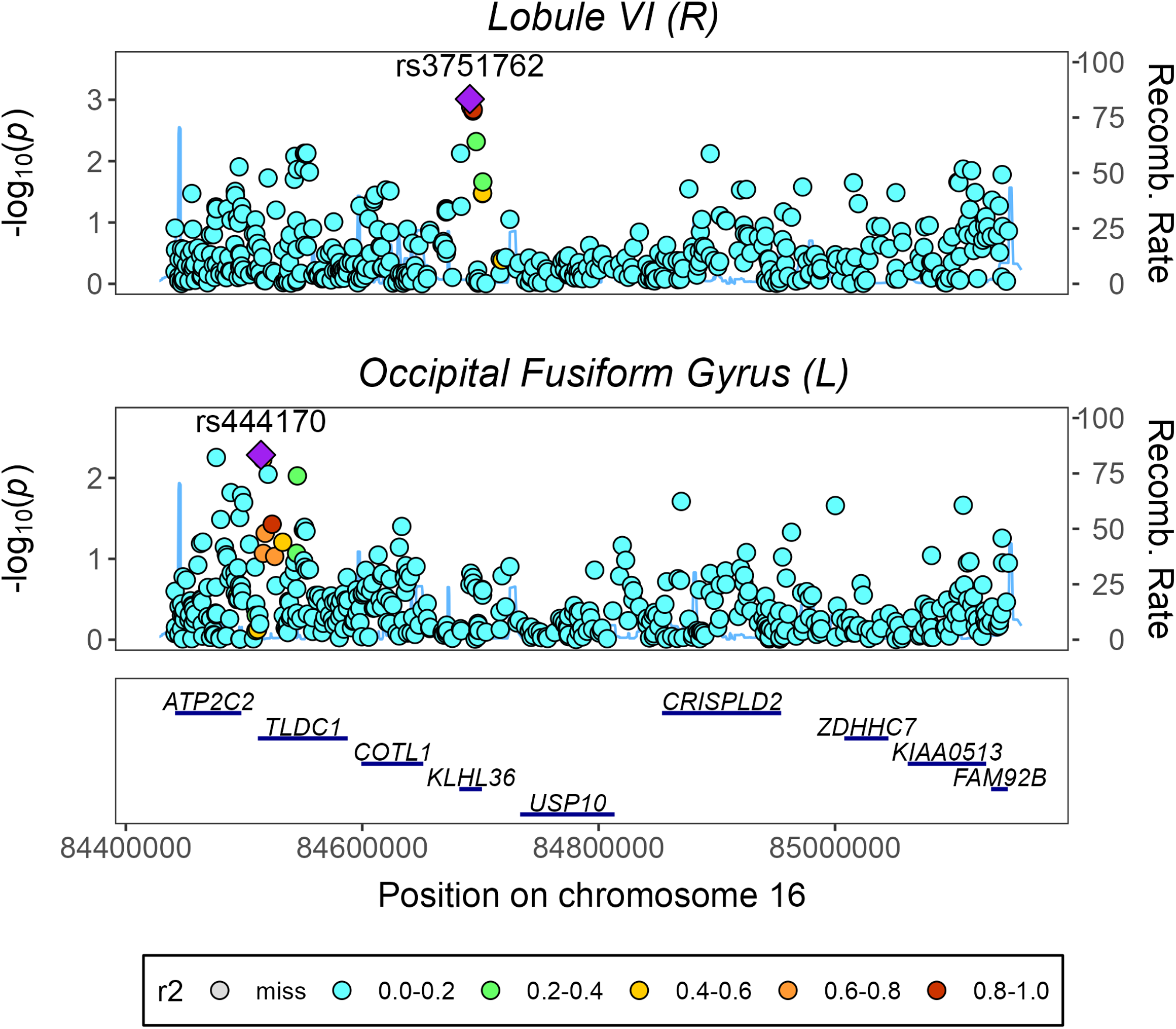
Regional association plot for traits with a significant local genomic correlation in locus 16:84440155-85146805, (LAVA). HapMap3 SNPs are shown.

**Figure S38.**
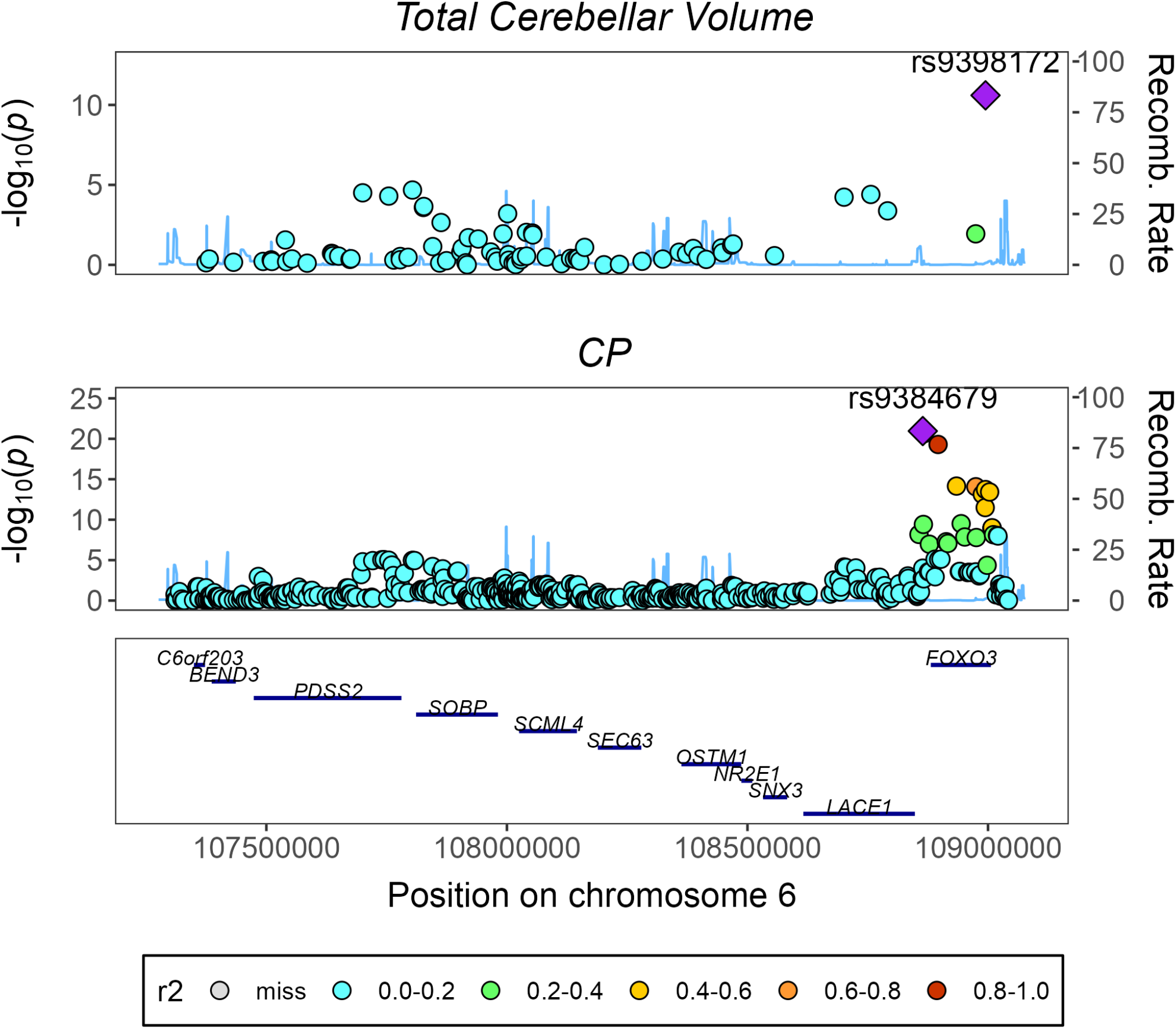
Regional association plot for traits with a significant local genomic correlation in locus 6:107309328-109043244, (LAVA). HapMap3 SNPs are shown.

**Figure S39.**
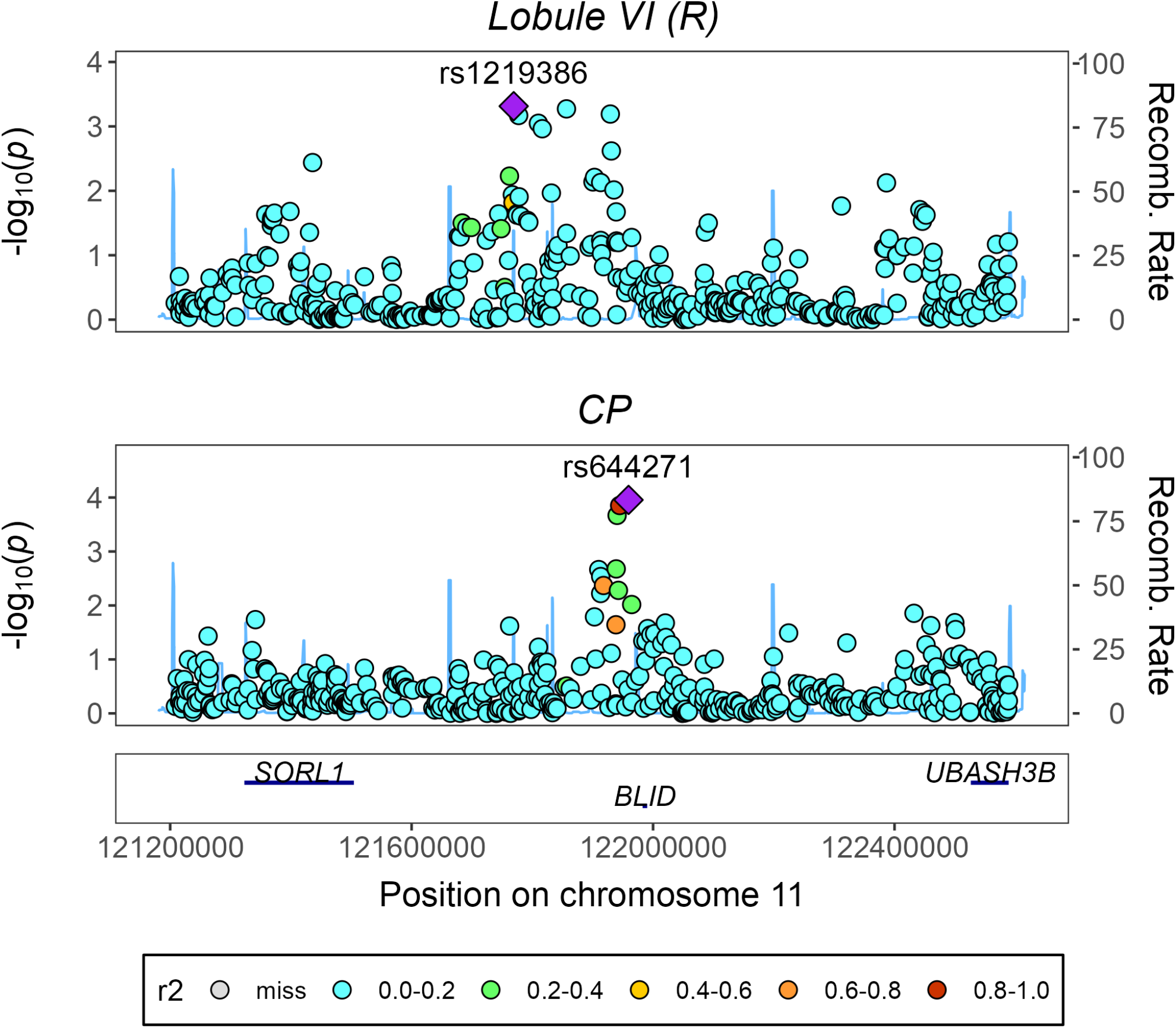
Regional association plot for traits with a significant local genomic correlation in locus 11:121208187-122589224, (LAVA). HapMap3 SNPs are shown.

**Figure S40.**
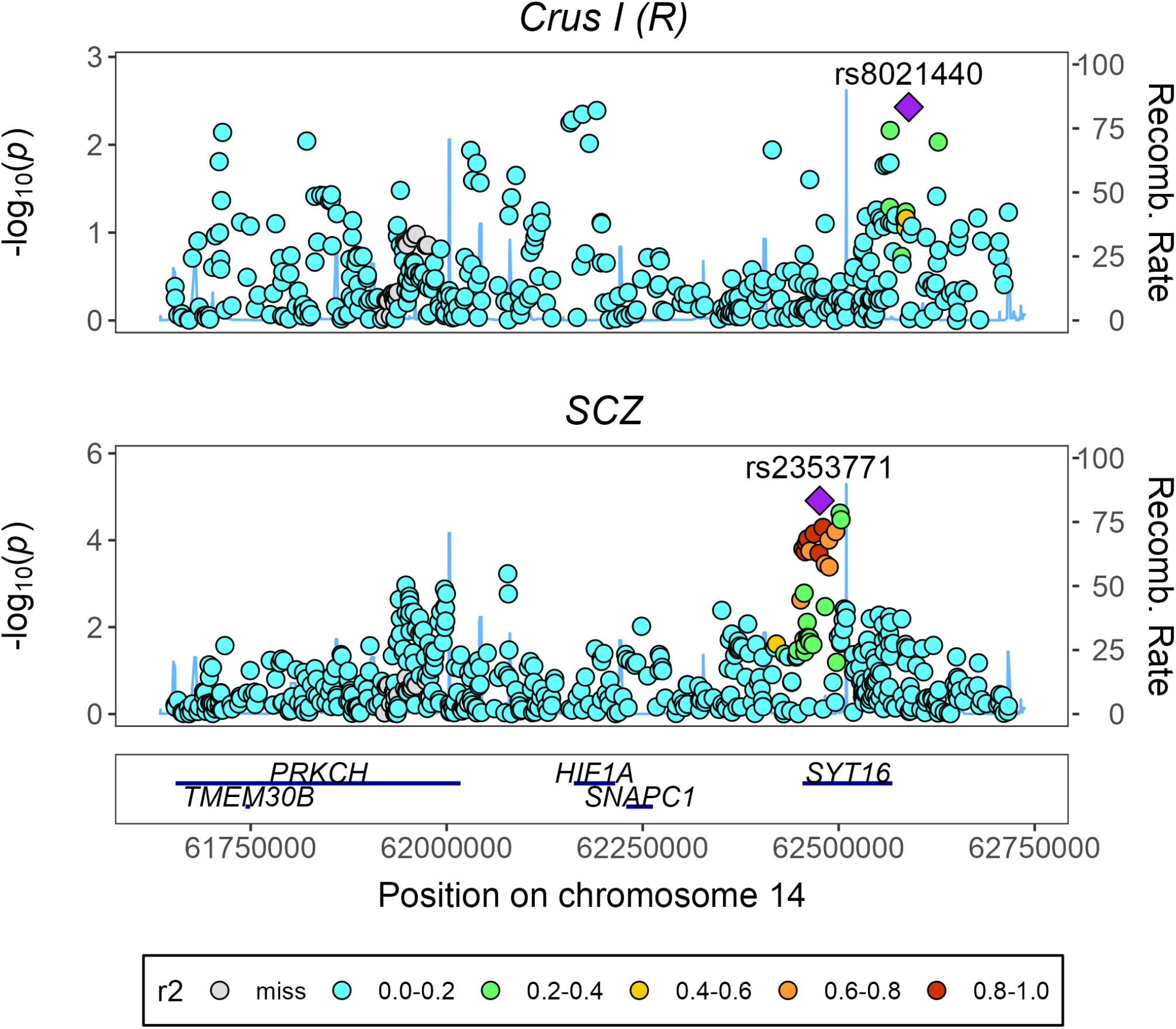
Regional association plot for traits with a significant local genomic correlation in locus 14:61652817-62717985, (LAVA). HapMap3 SNPs are shown.

**Figure S41.**
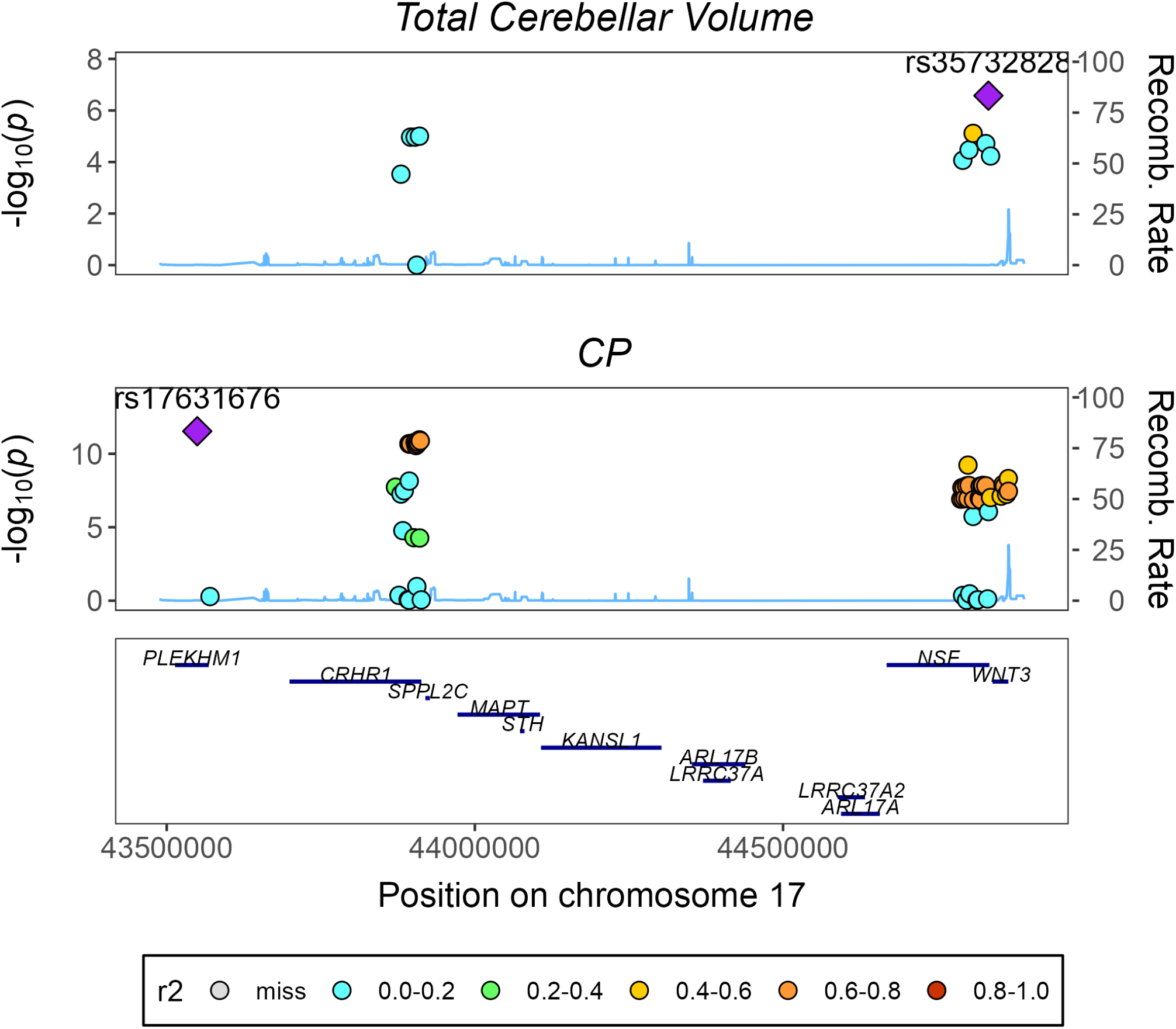
Regional association plot for traits with a significant local genomic correlation in locus 17:43460501-44865832, (LAVA). HapMap3 SNPs are shown.

